# Reprograming of the ubiquitin ligase Ubr1 by intrinsically disordered Roq1 through cooperating multifunctional motifs

**DOI:** 10.1101/2024.07.24.604893

**Authors:** Niklas Peters, Sibylle Kanngießer, Oliver Pajonk, Rafael Salazar Claros, Axel Mogk, Sebastian Schuck

## Abstract

One way cells control the speed and specificity of protein degradation is by regulating the activity of ubiquitin ligases. Upon proteotoxic stress in yeast, the intrinsically disordered protein Roq1 binds the ubiquitin ligase Ubr1 as a pseudosubstrate, thereby modulating the degradation of substrates of the N-degron pathway and promoting the elimination of misfolded proteins. The mechanism underlying this reprograming of Ubr1 is unknown. Here, we show that Roq1 controls Ubr1 by means of two cooperating multifunctional motifs. The N-terminal arginine and a short hydrophobic motif of Roq1 interact with Ubr1 as part of a heterobivalent binding mechanism. Via its N-terminal arginine, Roq1 regulates the ubiquitination of various N-degron substrates and folded proteins. Via its hydrophobic motif, Roq1 accelerates the ubiquitination of misfolded proteins. These findings reveal how a small, intrinsically disordered protein with a simple architecture engages parallel channels of communication to reprogram a functionally complex ubiquitin ligase.

## Introduction

The regulation of protein degradation is vital for cell homeostasis. By accelerating or slowing the degradation of certain proteins, cells can tune their activities to changing physiological demands. In addition, cells boost their capacity to degrade aberrant proteins when stress conditions provoke protein misfolding (McShane and Selbach, 2022). In eukaryotes, the degradation of thousands of proteins is carried out by the ubiquitin-proteasome system. Its selectivity is ensured by E3 ubiquitin ligases, which recognize degradation determinants (degrons) in target proteins and attach ubiquitin chains to these proteins as marks for destruction by the proteasome. The activities of ubiquitin ligases are controlled by a variety of mechanisms, such as phosphorylation, auto-ubiquitination and the interaction with protein or non-protein ligands (Vittal et al, 2015; Buetow and Huang, 2016; Zheng and Shabek, 2017). Unravelling these regulatory mechanisms is essential for understanding how cells orchestrate protein degradation.

A conserved and extensively studied ubiquitin ligase is Ubr1, which is best known for its central role in the N-degron pathway. It determines the half-life of proteins with specific N-terminal amino acid residues and thus controls a large number of processes, including chromosome segregation, inflammation, DNA repair and apoptosis (Varshavsky, 2011; Kim et al, 2021). Ubr1 consists of a single subunit that has a mass of about 200 kilodaltons and comprises a RING domain. The RING domain helps to bind a ubiquitin-loaded E2 ubiquitin-conjugating enzyme and enables the transfer of ubiquitin from the E2 enzyme onto lysine residues in substrate proteins (Deshaies and Joazeiro, 2009; Pan et al, 2021). Ubr1 recognizes N-degron substrates via two well-characterized binding sites, the type-1 site for proteins with positively charged N-terminal residues (type-1 N-degron substrates) and the type-2 site for proteins with bulky hydrophobic N-terminal residues (type-2 N-degron substrates). Furthermore, yeast Ubr1 contains a third, poorly-defined substrate binding site for folded proteins with internal degrons (Turner et al, 2000). These three binding sites communicate with each other so that occupancy of the type-1 and -2 sites allosterically and synergistically promotes substrate recognition by the third binding site (Du et al, 2002). Accordingly, the activity of Ubr1 towards certain endogenous proteins can be stimulated by peptides that bind to the type-1 or -2 sites, and this regulatory system controls peptide uptake in yeast (Turner et al, 2000). A similar system may counteract lipid droplet accumulation in hepatocytes (Zhang et al, 2022). Finally, Ubr1 can also recognize misfolded proteins, presumably through a fourth, yet uncharacterized substrate binding site (Eisele and Wolf, 2008; Heck et al, 2010).

In previous work in yeast, we discovered SHRED (stress-induced homeostatically regulated protein degradation), a regulatory cascade that is activated by proteotoxic stress, alters the substrate specificity of Ubr1 and thereby stimulates the degradation of misfolded proteins (Figure 1A; Szoradi et al, 2018). Specifically, various stress conditions that cause protein misfolding induce the synthesis of the small intrinsically disordered protein Roq1. Full-length Roq1 consists of 104 amino acid residues and is rapidly cleaved by the protease Ynm3. The resulting C-terminal cleavage fragment, Roq1(22-104), exposes arginine-22 (R22) at its new N-terminus. By means of the positively charged R22, Roq1(22-104) binds to the type-1 site of Ubr1 as a pseudosubstrate, enhances the ability of Ubr1 to eliminate misfolded proteins and thus increases cellular stress resistance. In addition, Roq1(22-104) competitively inhibits the degradation of type-1 substrates and promotes the degradation of type-2 substrates. Hence, SHRED involves comprehensive reprograming of Ubr1. However, beyond the interaction of Roq1 R22 and the Ubr1 type-1 site, the molecular mechanism of this unusual regulation remains unknown.

**Figure 1.**
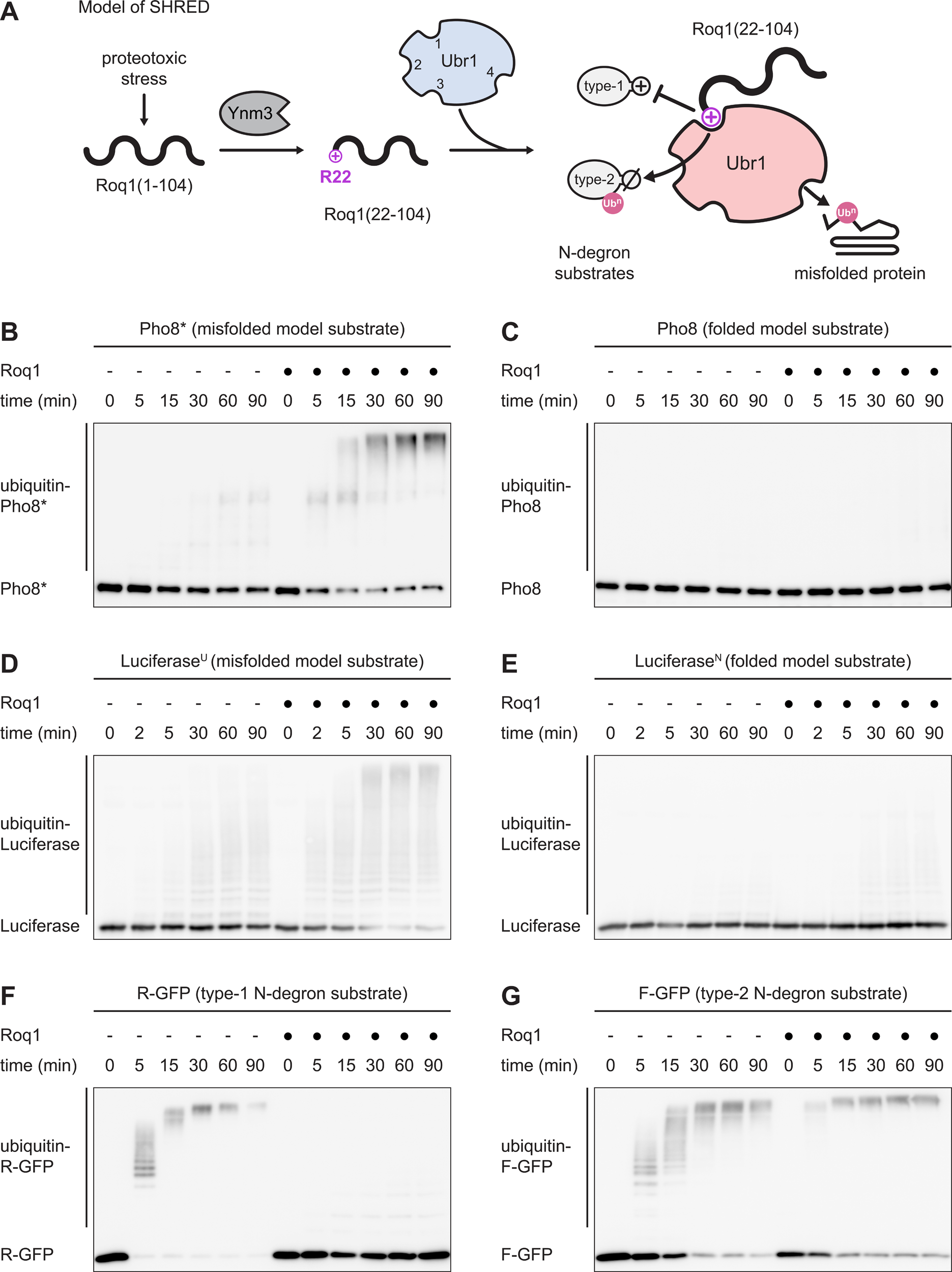
Roq1 reprograms Ubr1 substrate specificity in vitro. A Model of stress-induced homeostatically regulated protein degradation (SHRED). Proteotoxic stress in yeast induces the synthesis of Roq1(1-104), which is cleaved by the protease Ynm3 to generate Roq1(22-104) with arginine-22 (R22) at its new N-terminus. Through R22, Roq1(22-104) binds to the Ubr1 type-1 site as a pseudosubstrate and enhances the degradation of misfolded proteins to restore cell homeostasis. In addition, binding of Roq1(22-104) binding to the Ubr1 type-1 site competitively inhibits recognition of type-1 N-degron substrates and allosterically promotes recognition of type-2 N-degron substrates. Numbers in unbound Ubr1 (blue) denote the known and putative substrate binding sites for type-1 N-degron substrates (1), type-2 N-degron substrates (2), folded proteins with internal degrons (3) and misfolded proteins (4). Ub^n^, ubiquitin chain. B, C Western blot of Pho8 from Pho8* or Pho8 ubiquitination assays in the absence or presence of Roq1(22-60) for the times indicated. D, E Western blot of Luciferase from Luciferase^U^ or Luciferase^N^ ubiquitination assays in the absence or presence of Roq1(22-60) for the times indicated. F, G Western blot of GFP from R-GFP or F-GFP ubiquitination assays in the absence or presence of Roq1(22-60) for the times indicated.

In this study, we take an in vitro reconstitution approach to understand Ubr1 reprograming by Roq1. We show that Roq1 differentially modulates the ubiquitination of distinct types of Ubr1 substrates. We then define two cooperating sequence motifs in Roq1 that are needed for binding and regulating Ubr1. Finally, we provide evidence that these motifs engage parallel channels of communication between regulatory and substrate binding sites in Ubr1 to control its substrate specificity.

## Results

### In vitro reconstitution of Ubr1 reprograming by Roq1

To understand the mechanism of SHRED, we reconstituted the reprograming of Ubr1 by Roq1 in vitro. We showed previously that Roq1 cleavage by Ynm3 can be bypassed with Roq1(22-104), which lacks the first 21 residues of full-length Roq1 (Szoradi et al, 2018). We therefore used Roq1(22-104) for reconstitution experiments. To assay Ubr1 activity, we combined purified Roq1(22-104), ubiquitin, the ubiquitin-activating enzyme Ube1, ATP, the ubiquitin-conjugating enzyme Rad6, Ubr1 and potential Ubr1 substrates. However, we noticed that Ubr1 was also able to ubiquitinate Roq1(22-104), which, for our purposes, was an unwanted side reaction (Figure S1A; Pan et al, 2021). As Roq1(22-104) does not contain lysines, this ubiquitination likely occurs on serines, threonines or cysteines (Kelsall, 2022). The ubiquitin-Roq1 linkage was resistant to the dithiothreitol used to stop the ubiquitination reactions but was sensitive to mild alkaline hydrolysis (Figure S1A). Hence, the ubiquitin-Roq1 linkage must consist of oxyester rather than amide bonds. To focus our analysis on the activity of Ubr1 towards designated substrate proteins, we sought to eliminate the ubiquitination of Roq1. Shortening Roq1 to Roq1(22-60) almost completely eliminated ubiquitination (Figure S1A). Nevertheless, Roq1(22-60) was still able to support the stress-induced degradation of the SHRED reporter Rtn1-Pho8*-GFP in vivo (Figure S1B). We therefore mainly employed Roq1(22-60) for ubiquitination assays and Roq1(22-104) for interaction studies.

Using this simplified system, we tested several Ubr1 substrates. First, we analyzed the misfolded model protein Pho8*, which is a SHRED substrate in vivo (Szoradi et al, 2018). Ubr1 barely ubiquitinated Pho8* on its own but did so extensively in the presence of Roq1(22-60) (Figure 1B). By contrast, Ubr1 did not appreciably ubiquitinate Pho8, the folded counterpart of Pho8*, regardless of the absence or presence of Roq1 (Figure 1C). Control experiments showed that Pho8* remained mostly soluble throughout the assay, that its ubiquitination required Ubr1 and that ubiquitination occurred on lysines (Figure S1C-E). Roq1 already strongly promoted Pho8* ubiquitination when present at the same concentration as Ubr1, implying a tight interaction (Figure S1F). The amount of non-ubiquitinated Pho8* was reduced further with increasing Roq1 concentrations and we used a 10-fold molar excess of Roq1 over Ubr1 as a standard assay condition. Roq1(22-104) also enhanced Pho8* ubiquitination but less efficiently than Roq1(22-60) (Figure S1G). This can presumably be explained by the simultaneously occurring ubiquitination of Roq1(22-104), which diverts some Ubr1 from Pho8*. Next, we tested firefly Luciferase, a destabilized variant of which is a SHRED substrate in vivo (Szoradi et al, 2018). Roq1(22-60) stimulated Ubr1-mediated ubiquitination of unfolded Luciferase^U^ and had a much weaker effect on folded Luciferase^N^ (Figure 1D, E). These results confirm that Roq1 selectively promotes Ubr1-mediated ubiquitination of misfolded or unfolded proteins.

As a second type of substrate, we examined natural, well-folded proteins that are recognized by Ubr1 because of their internal degrons. These proteins were the transcriptional repressor Cup9, the DNA alkyltransferase Mgt1 and the mitotic checkpoint kinase Chk1 (Turner et al, 2000; Hwang et al, 2009; Oh et al, 2017). Roq1(22-60) stimulated Ubr1-mediated ubiquitination of all three proteins (Figure S2).

As a third type of substrate, we investigated proteins that are recognized by Ubr1 as part of the N-degron pathway. We showed previously that, in vivo, cleaved Roq1 binds to the Ubr1 type-1 site by means of its N-terminal R22 and thereby hampers the recognition of other proteins with positively charged N-terminal residues (Figure 1A; Szoradi et al, 2018). In agreement with these findings, in vitro ubiquitination of the type-1 N-degron model substrate R-GFP was inhibited by Roq1(22-60) (Figure 1F). Occupancy of the Ubr1 type-1 site facilitates the turnover of type-2 N-degron substrates with bulky hydrophobic N-terminal residues (Baker and Varshavsky, 1991). Accordingly, cleaved Roq1 promotes the in vivo turnover of N-degron substrates that can bind to the Ubr1 type-2 site (Szoradi et al, 2018). The same effect was apparent in vitro, where Roq1(22-60) enhanced Ubr1-mediated ubiquitination of the type-2 N-degron substrate F-GFP (Figure 1G). These observations corroborate that Roq1 inhibits the ubiquitination of type-1 N-degron substrates but promotes the ubiquitination of type-2 N-degron substrates.

Collectively, the in vitro reconstitution experiments demonstrated that Roq1 alters the substrate specificity of Ubr1: it promotes the ubiquitination of misfolded proteins, folded proteins with internal degrons and type-2 N-degron substrates, but inhibits the ubiquitination of type-1 N-degron substrates. Thus, the in vitro system contains all components minimally required for the reprograming of Ubr1 by Roq1.

### Roq1 contains a functionally essential hydrophobic motif

Next, we sought to determine how Roq1 reprograms Ubr1. We found previously that the N-terminal R22 of cleaved Roq1 is required for binding to the Ubr1 type-1 site (Figure 1A; Szoradi et al, 2018). Accordingly, an R22A mutant variant of Roq1(22-60) failed to stimulate Pho8* ubiquitination by Ubr1 (Figure 2A). A simple model for Roq1 action is that binding of R22 to the Ubr1 type-1 site allosterically activates the recognition of both type-2 N-degron substrates and misfolded proteins. To test this idea, we attempted to replace Roq1 with an arginine-alanine (RA) dipeptide, which binds to the Ubr1 type-1 site (Baker and Varshavsky, 1991). The RA dipeptide enhanced ubiquitination of the type-2 N-degron substrate F-GFP, as expected (Figure 2B). However, it did not stimulate the ubiquitination of Pho8*, even at a 4000-fold molar excess over Ubr1 (Figure 2C). Hence, occupancy of the Ubr1 type-1 site alone does not promote the ubiquitination of misfolded proteins.

**Figure 2.**
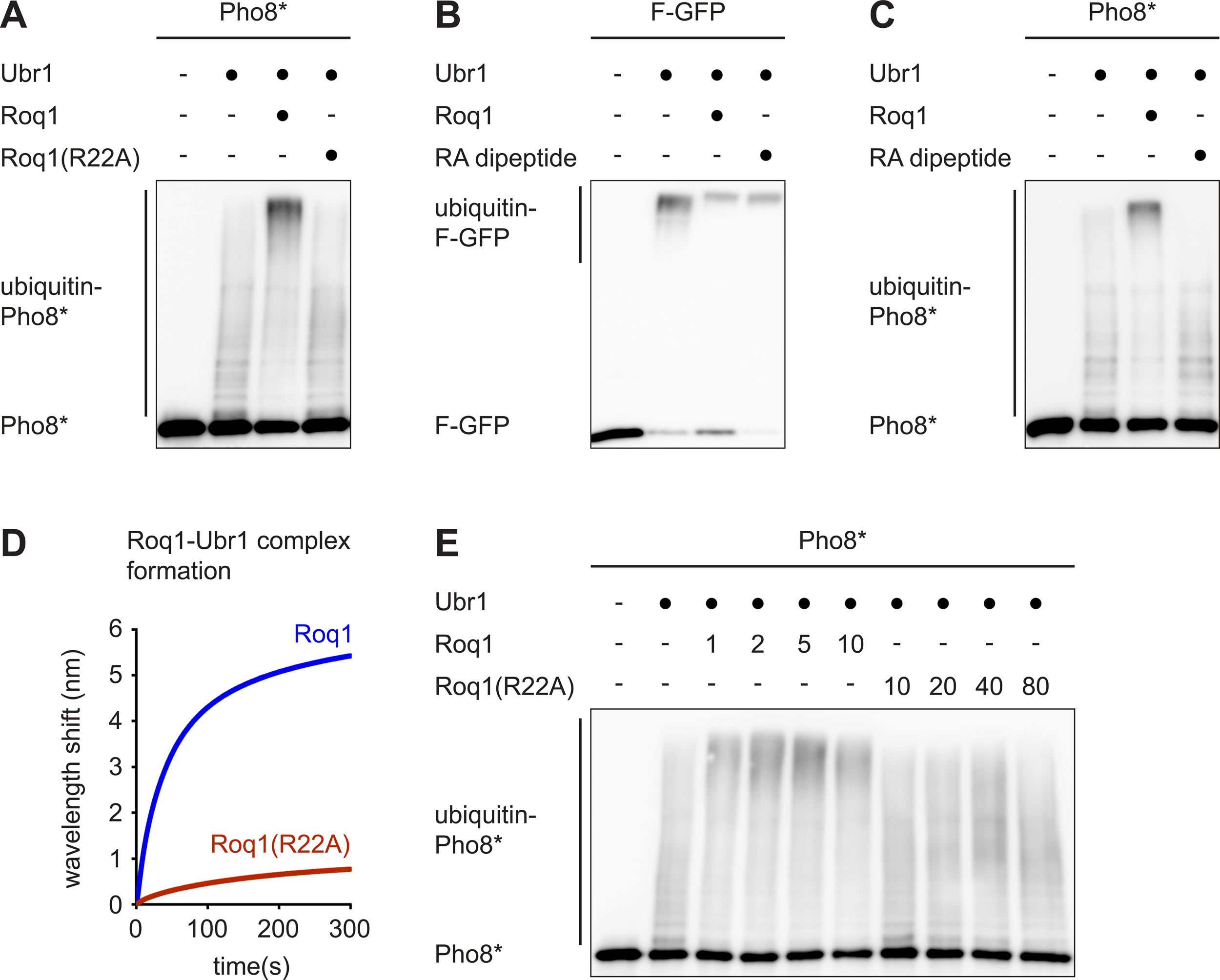
Roq1 binds and regulates Ubr1 via R22 and at least one other determinant. A Western blot of Pho8 from Pho8* ubiquitination assays with Roq1(22-60) or Roq1(22-60)(R22A). Roq1:Ubr1 molar ratios were 10:1. t = 90 min. B Western blot of GFP from F-GFP ubiquitination assays without and with Roq1(22-60) or the RA dipeptide. Molar ratios were Roq1:Ubr1 = 10:1 and RA:Ubr1 = 4000:1. t = 90 min. C As in panel B but western blot of Pho8 from Pho8* ubiquitination assays. D Biolayer interferometry of Roq1-Ubr1 complex formation with immobilized Roq1(22-104) and soluble Ubr1. Data are the mean of three independent experiments. E Western blot of Pho8 from Pho8* ubiquitination assays with Roq1(22-60) or Roq1(22-60)(R22A). Numbers in the Roq1 rows indicate the Roq1:Ubr1 molar ratios used. t = 90 min.

The data above imply that Roq1 harbors at least one determinant beyond R22 that is relevant for Ubr1 binding, Ubr1 regulation, or both. To study Roq1-Ubr1 interaction, we employed biolayer interferometry. We immobilized biotinylated Roq1(22-104) on a biosensor, added soluble Ubr1 and measured binding. Kinetic assays showed robust association of Ubr1 with Roq1. The association was diminished but not abolished by the R22A mutation, further confirming the presence of a second Ubr1 binding site in Roq1 (Figure 2D). Last, we asked whether Roq1 lacking R22 retained any capacity to activate Ubr1. As shown above, Roq1(R22A) did not affect Pho8* ubiquitination when used at a 10-fold molar excess over Ubr1 (Figure 2A). However, further raising its concentration slightly enhanced Pho8* ubiquitination (Figure 2E, for example note the upward shift of ubiquitin-Pho8* in the lane with a 40-fold molar excess of Roq1(R22A) compared with the lane with a 10-fold molar excess). Therefore, Roq1(R22A) still has a limited ability to activate Ubr1. These results show that cleaved Roq1 contains at minimum one determinant besides R22 that is important for both Ubr1 binding and regulation.

Roq1 is predicted to lack a defined tertiary structure and to be almost entirely disordered (Figure 3A, B; Jumper et al, 2021; Kurgan, 2022). In addition, sequence homologs of Roq1 can be recognized only in closely related fungi. Among fungal Roq1 homologs, the sequence around the Ynm3 cleavage site and a hydrophobic region in the middle of the protein sequence show the strongest conservation (Figure S3A). To identify functionally important features in Roq1 in an unbiased manner and potentially obtain informative Roq1 mutant variants, we carried out a genetic screen. As illustrated in Figure 3C, we mutagenized full-length Roq1 by error-prone PCR, introduced the resulting PCR products into *roq1* knockout yeast harboring the SHRED reporter Rtn1-Pho8*-GFP and seeded cells onto solid medium. SHRED-competent cells degrade Rtn1-Pho8*-GFP when they form mature colonies and lose their fluorescence, whereas SHRED-deficient cells remain fluorescent (Szoradi et al, 2018). To identify inactive Roq1 variants, we visually screened for colonies that retained fluorescence and sequenced their Roq1. We found fourteen unique point mutations, all of which impaired stress-induced Rtn1-Pho8*-GFP degradation as measured by flow cytometry (Figure 3D). In these experiments, Roq1 variants were expressed under the strong *GPD* promoter, and all mutant variants were detectable by western blotting (Figure S3B). In contrast, wild-type Roq1 expressed under the much weaker endogenous *ROQ1* promoter was not abundant enough for detection (Szoradi et al, 2018; also see Figure 4A, left panel). Thus, the mutant variants were present at levels that should, in principle, have been sufficient for Roq1 to fulfill its role in SHRED. We therefore conclude that the mutations do not simply destabilize Roq1 but genuinely disrupt its function.

**Figure 3.**
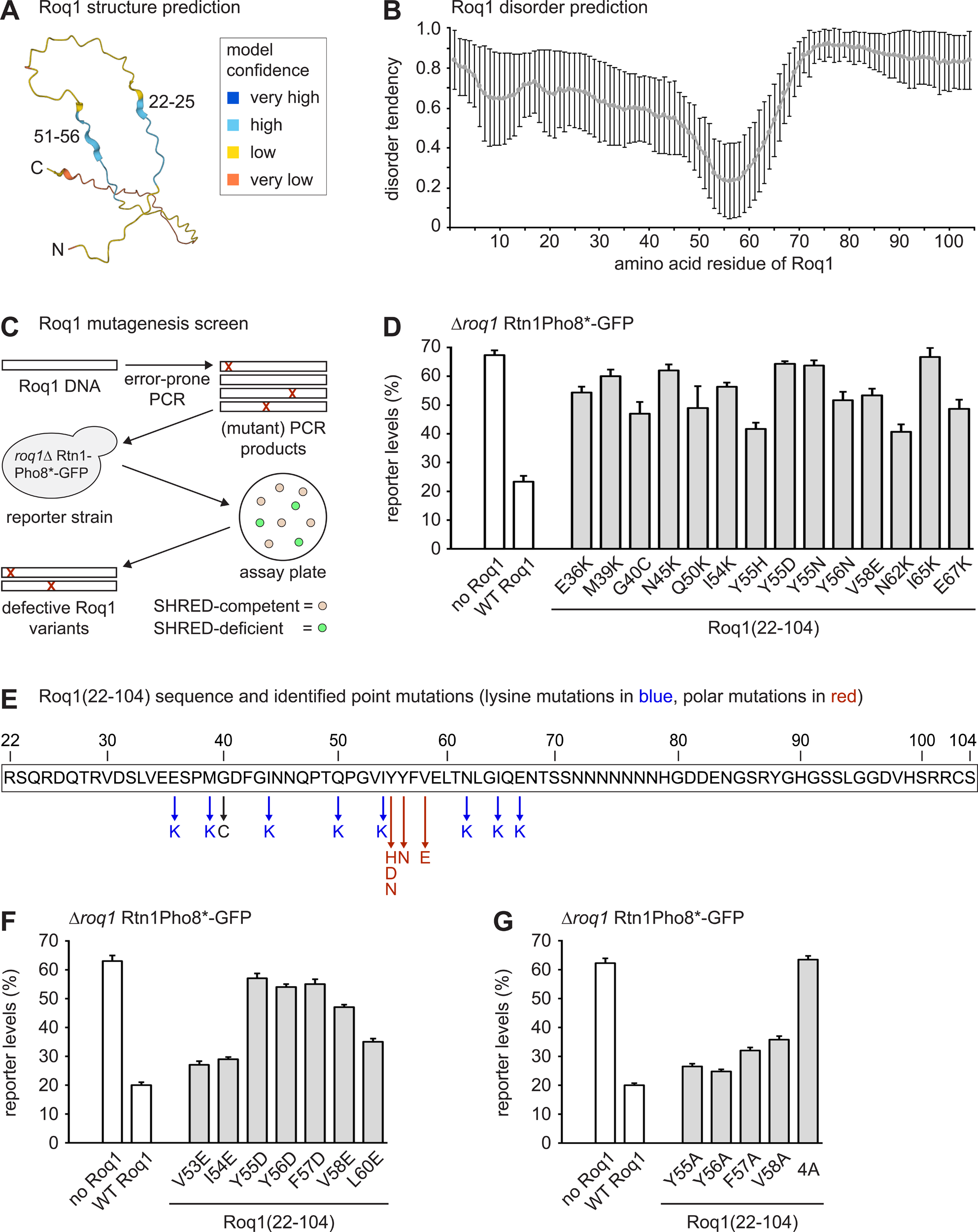
Roq1 contains a functionally essential hydrophobic motif. A Roq1 structure prediction by AlphaFold. B Roq1 disorder prediction. The plot shows the average disorder tendency across the Roq1(1-104) sequence on a scale of 0 to 1, as given by twelve different algorithms for disorder prediction. Error bars show the standard deviation. C Workflow of the Roq1 mutagenesis screen. Roq1 variants mutagenized by error-prone PCR were introduced into a reporter strain lacking endogenous Roq1. Cells were allowed to form colonies on assay plates, SHRED-deficient colonies were picked and their defective Roq1 variants were sequenced. D Cellular levels of the SHRED reporter Rtn1-Pho8*-GFP after tunicamycin treatment for 5 h relative to levels in untreated cells, as measured by flow cytometry. The *roq1* mutant cells contained an empty plasmid (no Roq1), a plasmid encoding wild-type Roq1 (WT Roq1) or plasmids encoding variants of ubiquitin-Roq1(22-104) fusions. Ubiquitin-Roq1 fusions are processed by cells to yield Roq1(22-104). Bars are the mean ± SEM; n = 3 biological replicates. E Schematic of the Roq1(22-104) sequence showing the point mutations identified in the Roq1 mutagenesis screen. F, G As in panel D. 4A = Y55A,Y56A,F57A,V58A.

**Figure 4.**
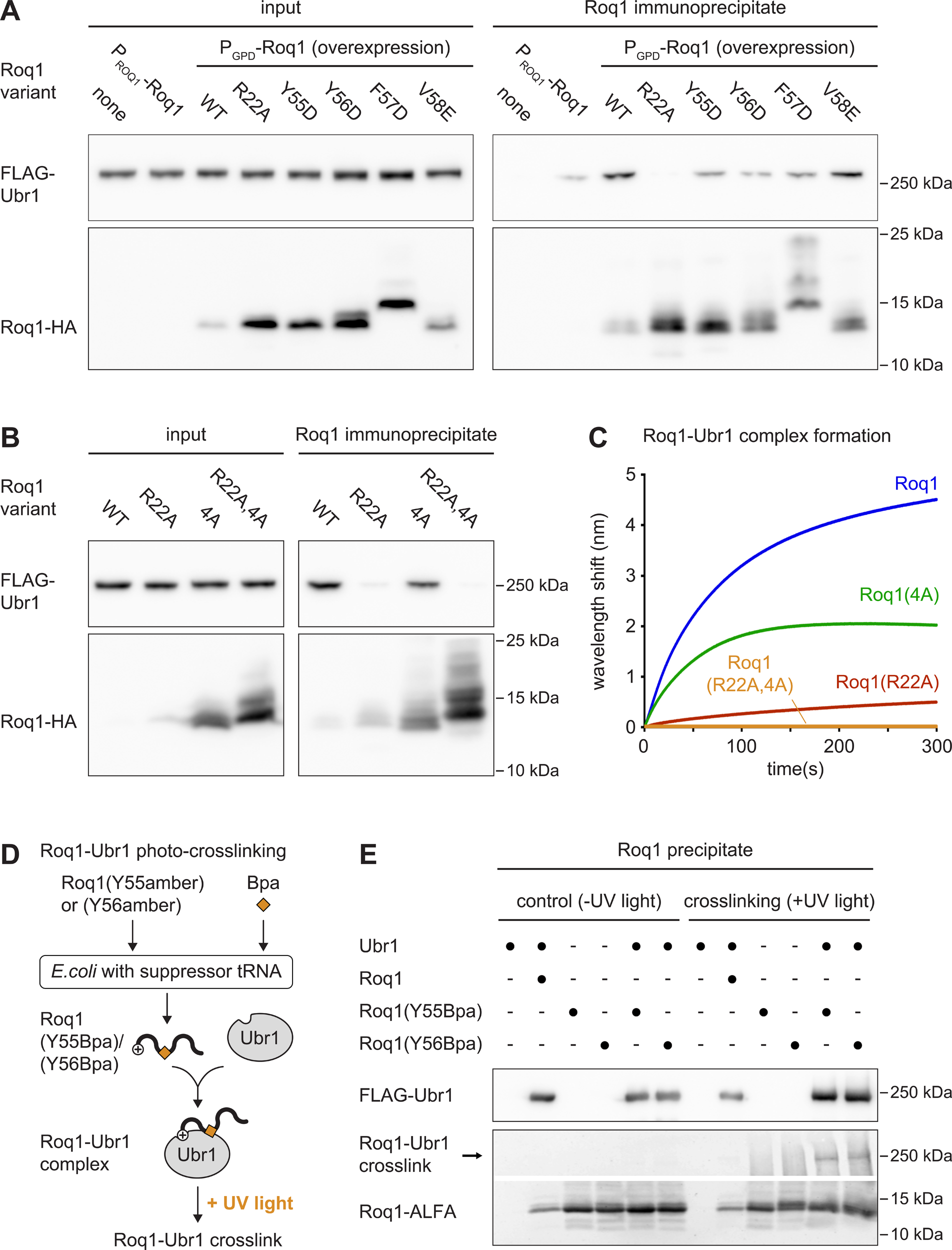
Roq1 interacts with Ubr1 through a heterobivalent binding mechanism. A Western blot of FLAG and HA tags from cell lysates (input) or anti-HA immunoprecipitates of *roq1* knockout cells expressing FLAG-Ubr1 and HA-tagged Roq1 variants as indicated (Roq1 immunoprecipitate). All Roq1 variants were expressed as ubiquitin-Roq1(22-104)-HA(74) fusions, which are processed by cells to yield Roq1(22-104)-HA(74) with an internal HA tag. Note that Roq1(22-104)-HA(74) expressed under the endogenous *ROQ1* promoter is undetectable due to low expression levels. Roq1(22-104) naturally runs as a double band. The nature of the slower migrating band is unknown, as is the reason for the strong impact of the F57D mutation on Roq1 electrophoretic mobility. P_ROQ1_, *ROQ1* promoter; P_GPD_, *GPD* promoter. B As in panel A. All Roq1 variants were expressed under the strong *GPD* promoter. C Biolayer interferometry of Roq1-Ubr1 complex formation with immobilized Roq1(22-104) and soluble Ubr1. Data are the mean of two independent experiments. 4A = Y55A,Y56A,F57A,V58A. D Workflow Roq1-Ubr1 photo-crosslinking. Roq1 with an amber stop codon instead of Y55 or Y56 was expressed in *E. coli* equipped to suppress the amber stop codon by incorporation of p-benzoyl-L-phenylalanine (Bpa). Roq1 was purified and mixed with Ubr1 to reconstitute the Roq1-Ubr1 complex. The complex was precipitated, irradiated with UV light and analyzed by western blotting. E Western blot of FLAG and ALFA tags from precipitated Roq1-Ubr1 complex with and without photo-crosslinking with UV light.

The mutations were exclusively in the Roq1(22-104) fragment and mostly fell into one of two classes that we termed lysine mutations and polar mutations (Figure 3E). Lysine mutations introduced lysine residues into the otherwise lysine-free Roq1(22-104) and were scattered across the Roq1 sequence. These mutations may inactivate Roq1 because they help Ubr1 to ubiquitinate Roq1 and thereby prevent Roq1-Ubr1 interaction. Consistent with this interpretation, changing residue I54 to an arginine instead of a lysine yielded active Roq1 (Figure S3C). This observation suggests that, at least for this residue, the lysine mutation disrupts Roq1 function by allowing Roq1 ubiquitination, rather than by introducing a positive charge. Furthermore, individually changing all residues affected by lysine mutations to alanines did not perturb the SHRED activity of Roq1 (Figure S3D). Hence, these residues are not critical for Roq1 function and we did not analyze the lysine mutations further. The polar mutations introduced polar or charged residues in place of hydrophobic residues clustered at positions 55-58. This region is the least likely part of Roq1 to be disordered and also the most hydrophobic (Figure 3B, E). We therefore individually mutated each hydrophobic residue from V53 to L60 to negatively charged residues. This analysis defined the Y55-Y56-F57-V58 sequence as critical for Roq1 function (Figure 3F). Individual mutation of these residues to alanines did not impair Ubr1 regulation by Roq1, but simultaneous mutation of all four residues to alanines abolished it (Figure 3G). Hence, the YYFV sequence, which we call the hydrophobic motif, is a second feature of Roq1 that is essential for SHRED.

Of note, mutations in the hydrophobic motif increased the levels of Roq1(22-104), indicating that the hydrophobic motif destabilizes Roq1 in vivo and may serve as a degron (Figure S3B). Roq1 is short-lived but Ubr1 has only a minor role in the turnover of wild-type Roq1 (Szoradi et al, 2018). Hence, Roq1 degradation must involve other ubiquitin ligases or occur in a ubiquitin-independent manner. In any event, it appears that the hydrophobic motif is both a degron and a critical determinant for Ubr1 regulation.

### Roq1 interacts with Ubr1 through a heterobivalent binding mechanism

Next, we asked whether the hydrophobic motif was required for the interaction of Roq1 and Ubr1. We expressed HA-tagged wild-type and mutant variants of Roq1(22-104) in yeast harboring FLAG-Ubr1, immunoprecipitated Roq1 and probed for co-precipitation of Ubr1. In agreement with earlier findings (Szoradi et al, 2018), wild-type Roq1 efficiently precipitated Ubr1, and this interaction was strongly diminished by the R22A mutation (Figure 4A). Point mutations that introduced negative charges into the hydrophobic motif reduced the affinity of Roq1 to Ubr1 to varying degrees. Mutation of all four residues of the hydrophobic motif to alanines reduced binding to Ubr1, and combined mutation of R22 and the hydrophobic motif abolished it (Figure 4B).

An interpretation of these in vivo experiments is that Roq1 interacts with Ubr1 through a heterobivalent binding mechanism that involves binding of R22 to the Ubr1 type-1 site and binding of the hydrophobic motif to another region of Ubr1. To test this model in vitro, we performed biolayer interferometry with Ubr1 and Roq1 variants. Individual mutation of R22 or the hydrophobic motif impaired Roq1-Ubr1 binding, and the combined mutation of R22 and the hydrophobic motif rendered the interaction undetectable (Figure 4C). To determine whether the hydrophobic motif was only indirectly involved in Roq1-Ubr1 interaction or directly bound to Ubr1, we employed photo-crosslinking. As illustrated in Figure 4D, we replaced Y55 or Y56 within the hydrophobic motif of Roq1(22-104) with an amber stop codon. We then purified these Roq1 variants from *E. coli* that suppress amber stop codons by incorporation of the non-natural, photo-crosslinkable amino acid p-benzoyl-L-phenylalanine (Bpa; Chin et al, 2002). Wild-type Roq1 and Bpa-containing variants were used to reconstitute the Roq1-Ubr1 complex, irradiated with UV light to induce crosslinks and analyzed by western blotting. Upon crosslinking, a fraction of Bpa-containing but not of wild-type Roq1(22-104) shifted from an apparent molecular mass of less than 15 kDa to roughly 250 kDa (Figure 4E). This apparent molecular mass corresponded to that of Ubr1, showing that Roq1 can be crosslinked to Ubr1. Hence, the Roq1 hydrophobic motif is likely to make direct physical contact with Ubr1.

In summary, these results demonstrate synergy between two points of physical contact between Roq1 and Ubr1 as part of a heterobivalent binding mechanism. One contact is provided by R22 of cleaved Roq1 and the Ubr1 type-1 site. The other contact involves the Roq1 hydrophobic motif and a site in Ubr1 that remains to be determined.

### The Roq1 hydrophobic motif selectively promotes the ubiquitination of misfolded proteins

We next wanted to determine whether the hydrophobic motif merely served as a binding determinant or, like R22, additionally controlled Ubr1 activity. For this purpose, we took advantage of the V58E mutation identified in the genetic screen. The V58E mutation crippled the ability of Roq1 to drive degradation of the SHRED reporter Rtn1-Pho8*-GFP in yeast (Figure 3D). Nonetheless, Roq1(V58E) still co-immunoprecipitated Ubr1 from cell lysates almost as efficiently as wild-type Roq1 (Figure 4A). Biolayer interferometry confirmed that Roq1(V58E) was still able to associate with Ubr1, albeit less rapidly than wild-type Roq1 (Figure 5A). These observations raised the possibility that the hydrophobic motif has separable binding and regulatory roles. Specifically, the hydrophobic motif could (1) stabilize the Roq1-Ubr1 interaction in a manner that involves but does not strictly depend on V58 and (2) activate Ubr1 in a manner that requires V58. To explore this idea, we compared the impact of wild-type Roq1 and Roq1(V58E) on the ubiquitination of different Ubr1 substrates. Roq1(V58E) could not stimulate Ubr1-mediated ubiquitination of Pho8* and only mildly enhanced ubiquitination of unfolded Luciferase^U^, even when the concentration of Roq1(V58E) was raised to compensate for its reduced binding to Ubr1 (Figure 5B, C). In striking contrast, wild-type Roq1 and Roq1(V58E) were equally effective at inhibiting ubiquitination of the type-1 N-degron substrate R-GFP, at stimulating ubiquitination of the type-2 N-degron substrate F-GFP and at stimulating ubiquitination of the folded Ubr1 substrate Cup9 (Figure 5D-F). We conclude that the key event in the control of R-GFP, F-GFP and Cup9 ubiquitination is binding of the Roq1 N-terminus to the Ubr1 type-1 site. For misfolded Pho8* and unfolded Luciferase^U^, however, the hydrophobic motif provides an additional regulatory activity that depends on V58.

**Figure 5.**
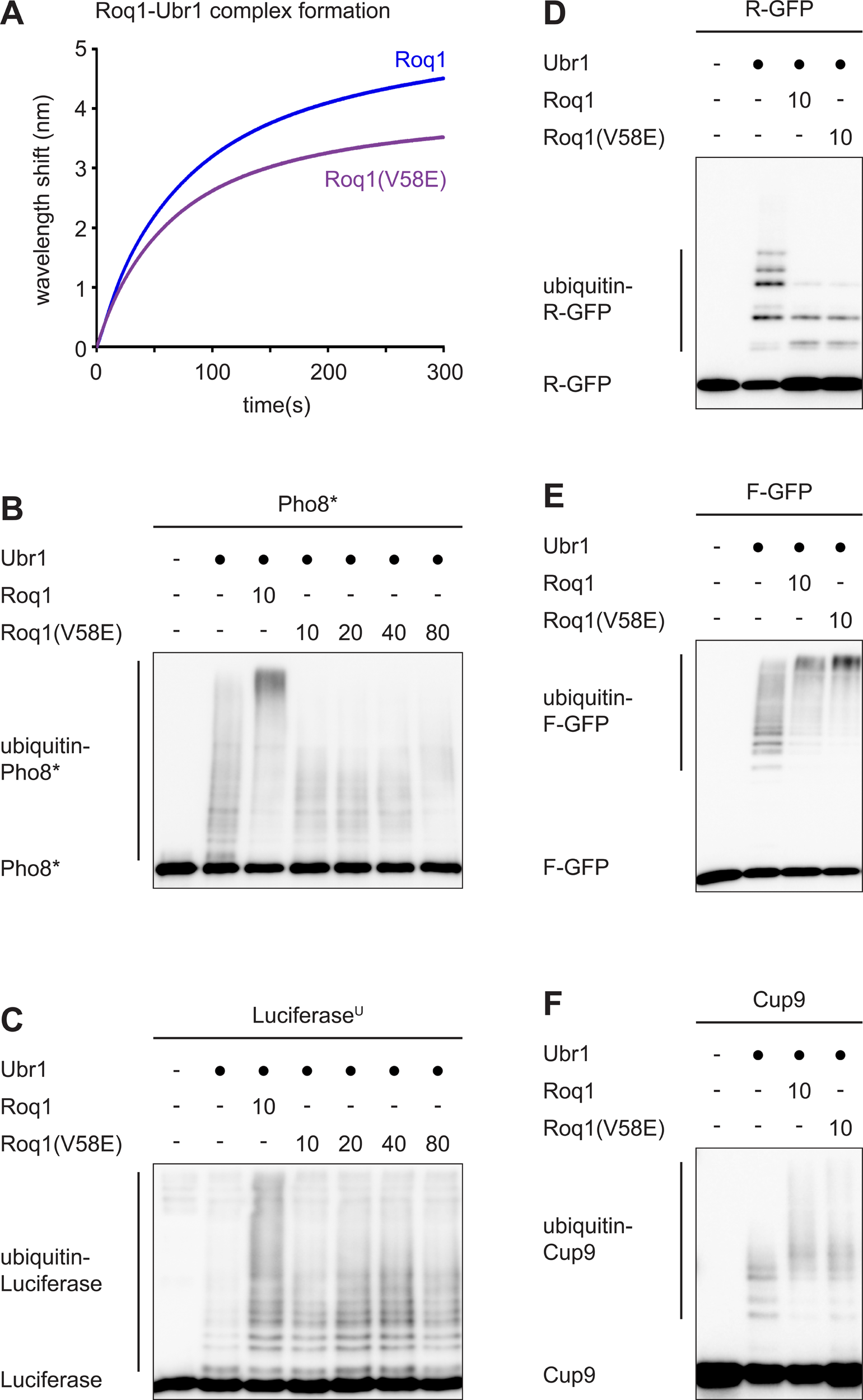
The Roq1 hydrophobic motif selectively promotes the ubiquitination of misfolded proteins. A Biolayer interferometry of Roq1-Ubr1 complex formation with immobilized Roq1(22-104) and soluble Ubr1. Data are the mean of two independent experiments. The data for wild-type Roq1 are the same as in Figure 4C. B Western blot of Pho8 from Pho8* ubiquitination assays with Roq1(22-60) or Roq1(22-60)(V58E). Numbers in the Roq1 rows indicate the Roq1:Ubr1 molar ratios used. t = 90 min. C As in panel B but western blot of Luciferase from Luciferase^U^ assays. t = 30 min. D As in panel B but western blot of GFP from R-GFP assays. t = 15 min. E As in panel B but western blot of GFP from F-GFP assays. t = 15 min. F As in panel B but western blot of Strep tag from Cup9-Strep assays. t = 15 min.

The Roq1 hydrophobic motif could regulate Ubr1 in several ways. It could change the conformation of Ubr1 to enhance the recognition of misfolded proteins, stimulate the recruitment of the ubiquitin-conjugating enzyme Rad6 or alter the structure of the Rad6-ubiquitin-Ubr1-substrate complex such that substrate ubiquitination becomes more efficient. To test these possibilities, we first used an in vitro pulldown assay to determine whether Roq1 affected the recognition of Pho8 or Pho8* by Ubr1. Pho8* bound to Ubr1 more strongly than Pho8, but Roq1 did not enhance this interaction (Figure S4A). Second, we analyzed the interaction of Rad6 and Ubr1 by biolayer interferometry. Rad6 can bind to Ubr1 even when not conjugated to ubiquitin (Xie and Varshavsky, 1999), and the association of unloaded Rad6 and Ubr1 was indeed detectable. However, Roq1 did not enhance it (Figure S4B). Third, we asked how the Roq1 hydrophobic motif stimulates substrate ubiquitination. Standard ubiquitination assays cannot distinguish whether Roq1 promotes chain initiation, i.e. the attachment of the first ubiquitin to lysines in substrate proteins, makes chain elongation more efficient, or has a combination of effects. To focus on chain initiation, we employed lysine-free ubiquitin (no-K-ubiquitin), which can be attached to substrate proteins via its C-terminus but cannot form ubiquitin chains. When we used no-K-ubiquitin together with Pho8*, Ubr1 on its own was able to generate two discrete bands that corresponded to Pho8* conjugated with one or two molecules of no-K-ubiquitin (Figure 6A). Hence, without Roq1, two different lysines of Pho8* can be efficiently ubiquitinated. Wild-type Roq1 increased the number of ubiquitinated lysines to at least five. Roq1(V58E) and the RA dipeptide had no effect, consistent with the absence of the regulatory activity provided by the hydrophobic motif. For F-GFP, Ubr1 on its own could attach ubiquitin to five distinct lysines (Figure 6B). The RA dipeptide raised the number of ubiquitinated lysines to six and increased the efficiency of ubiquitination. As expected, Roq1 and Roq1(V58E) had no effects beyond that of the RA dipeptide. The same was true for Cup9 (Figure 6C).

**Figure 6.**
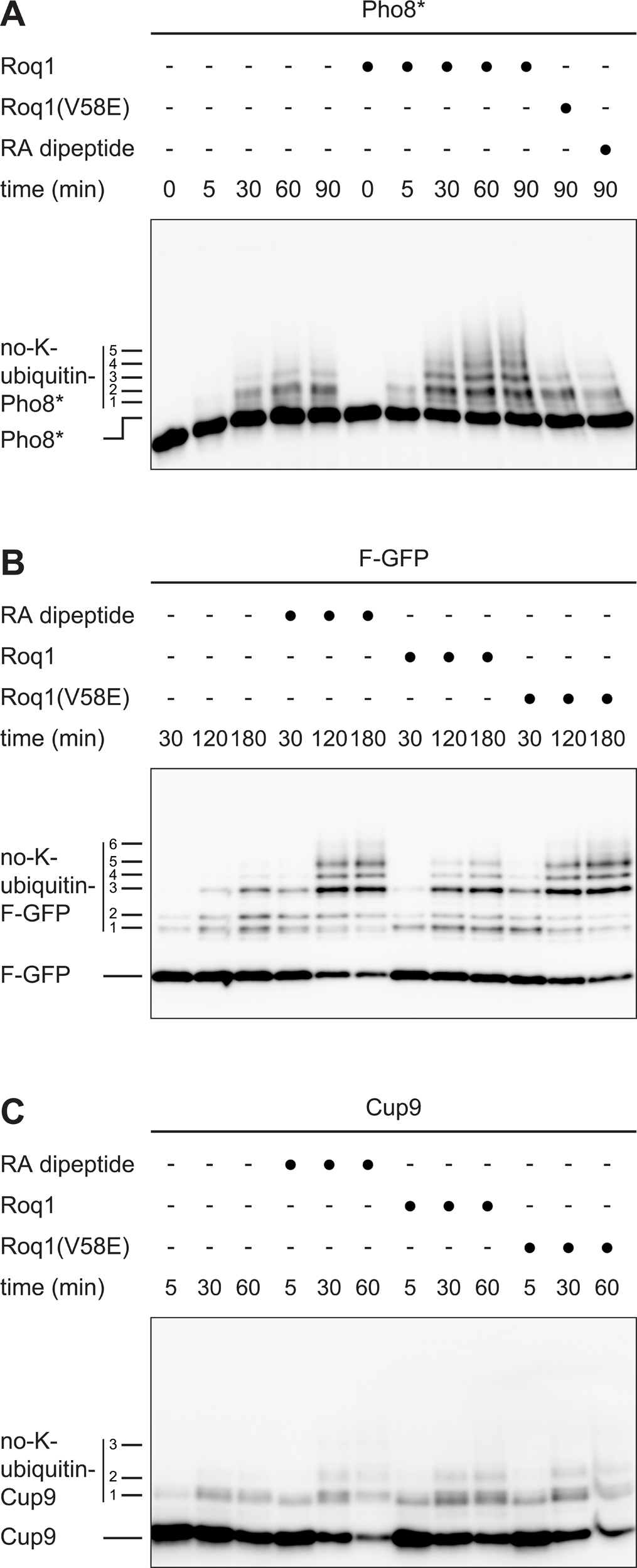
Roq1 promotes ubiquitin chain initiation on Ubr1 substrate proteins. A Western blot of Pho8 from Pho8* ubiquitination assays with lysine-free ubiquitin (no-K-ubiquitin) and Roq1(22-60), Roq1(22-60)(V58E) or RA dipeptide. Numbers to the left of the gel indicate distinct conjugates of no-K-ubiquitin and Pho8*. B As in panel A but western blot of GFP from F-GFP assays. C As in panel A but western blot of Strep tag from Cup9-Strep assays.

In summary, the Roq1 hydrophobic motif appears to have two functions. First, it stabilizes the association of Roq1 and Ubr1, which reinforces binding of R22 to the Ubr1 type-1 site. In this way, the hydrophobic motif helps R22 to regulate the ubiquitination of type-1 and type-2 N-degron substrates and folded substrates with internal degrons, such as Cup9. Second, the hydrophobic motif promotes the ubiquitination of misfolded proteins, at least in part by enhancing chain initiation. This second function specifically requires V58.

### Functional Roq1 minimally consists of R22, a short linker and the hydrophobic motif

To understand the functional architecture of Roq1 more completely, we aimed to reduce it to its minimally required parts. The data so far showed that Roq1 functionality in vitro and in vivo requires R22 and the hydrophobic motif. Therefore, we tested directly whether regions outside of these determinants were dispensable. As shown above, Roq1(22-104) and also Roq1(22-60) rescued SHRED in *roq1* knockout cells when expressed under the strong *GPD* promoter (Figure S1B). By contrast, Roq1(22-104) also rescued SHRED when expressed under the weak *ROQ1* promoter but Roq1(22-60) did so only partially (Figure S5). A possible explanation is that the sequence beyond residue 60 contributes to Roq1 stability in vivo. Nonetheless, the data with overexpressed Roq1(22-60) show that this sequence does not contain functionally essential determinants.

Next, we shortened the sequence that links R22 and the hydrophobic motif, which naturally consists of 33 amino acids (Figure 7A). One Roq1 variant that still restored substantial SHRED activity in *roq1* knockout cells upon expression under the *GPD* promoter was Roq1(22 aa) (Figure 7B). In this variant, only 15 residues between R22 and the hydrophobic motif remained. To test whether this sequence harbors functionally important residues, we replaced most residues with generic GGS- or GSP-based linkers (Figure 7C). Roq1(GGS) and Roq1(GSP) were inactive in vivo, possibly due to low stability (not shown). However, Roq1(22 aa), Roq1(GGS) and Roq1(GSP) were all fully active in vitro (Figure 7D). These experiments define minimalistic Roq1-derived peptides that are able to regulate Ubr1. Remarkably, peptides of only 22 residues that are composed of an N-terminal arginine residue, a flexible linker and a short hydrophobic motif is all that is needed to profoundly change the activity and substrate specificity of a large ubiquitin ligase that consists of nearly 2000 residues.

**Figure 7.**
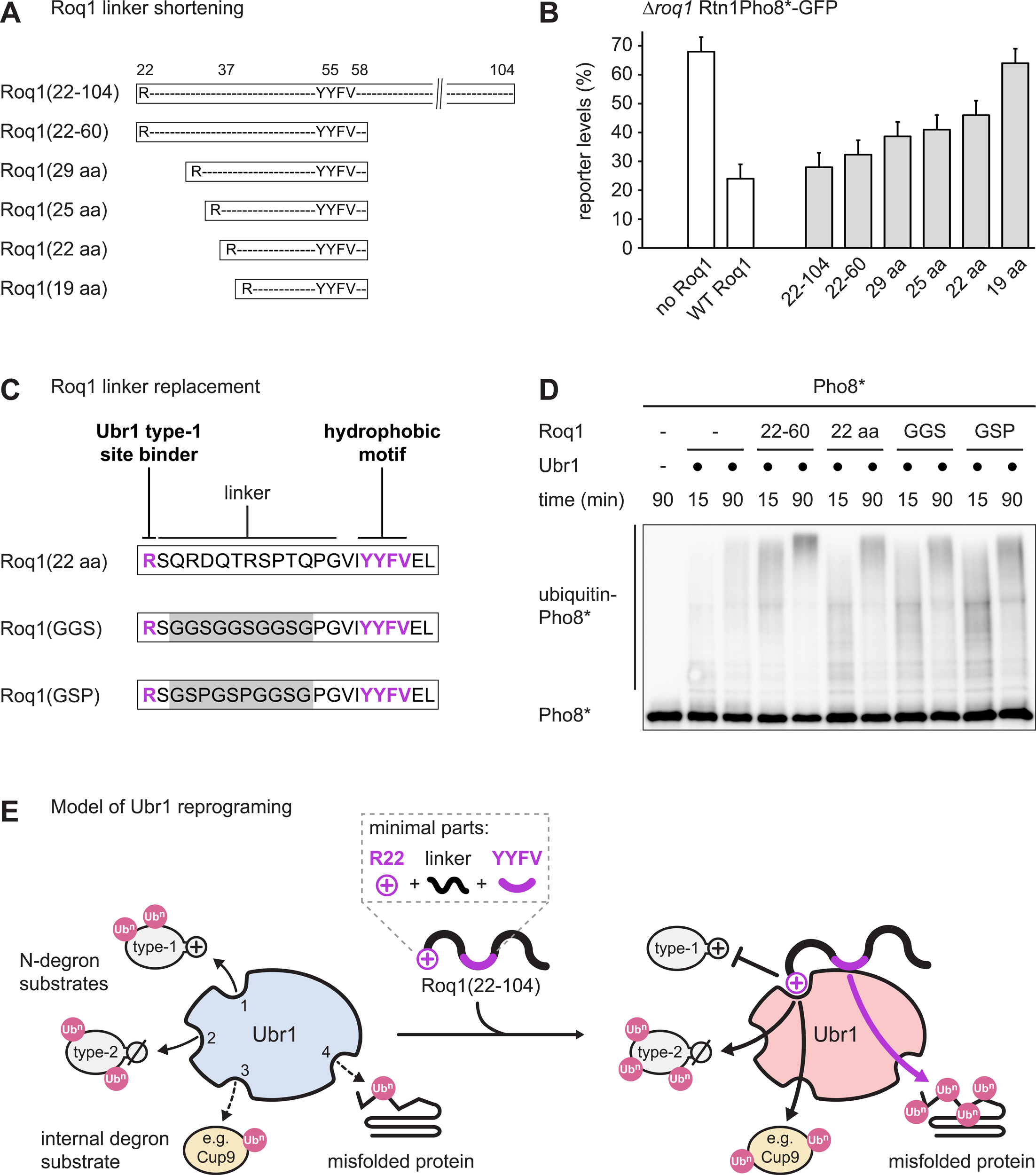
Functional Roq1 minimally consists of R22, a short linker and the hydrophobic motif. A Schematic of Roq1 linker shortening. B Cellular levels of the SHRED reporter Rtn1-Pho8*-GFP after tunicamycin treatment for 5 h relative to levels in untreated cells, as measured by flow cytometry. The *roq1* mutant cells contained an empty plasmid (no Roq1), a plasmid encoding wild-type Roq1 (WT Roq1) or plasmids encoding variants of ubiquitin-Roq1 fusions of various lengths. Ubiquitin-Roq1 fusions are processed by cells to yield Roq1 starting with R22. Bars are the mean ± SEM; n = 3 technical replicates. C Schematic of Roq1 constructs for replacement of the linker between R22 and the hydrophobic motif. D Western blot of Pho8 from Pho8* ubiquitination assays with Roq1 variants with shortened or artificial linkers between R22 and the hydrophobic motif. E Model of Ubr1 reprograming by Roq1. Cleaved Roq1(22-104) interacts with Ubr1 through a heterobivalent binding mechanism that creates avidity. Binding involves physical contact of R22 with the Ubr1 type-1 site and of the YYFV hydrophobic motif with an undefined region in Ubr1. Occupancy of the Ubr1 type-1 site activates the type-2 substrate binding site and a binding site for folded substrates with internal degrons, such as Cup9. In parallel, the hydrophobic motif activates a putative fourth substrate binding site to enhance the degradation of misfolded proteins. Numbers in unbound Ubr1 (blue) denote the known and putative substrate binding sites for type-1 N-degron substrates (1), type-2 N-degron substrates (2), folded proteins with internal degrons (3) and misfolded proteins (4). The box above Roq1(22-104) highlights the minimally required functional elements of Roq1. Ub^n^, ubiquitin chain.

## Discussion

Our findings reveal that Roq1 reprograms Ubr1 by simultaneously engaging two channels of communication. Roq1 uses R22 to occupy the Ubr1 type-1 site as a pseudosubstrate. This interaction activates the Ubr1 type-2 site and the binding site for folded substrates with internal degrons. In parallel, Roq1 uses the hydrophobic motif to bind to Ubr1 at a separate regulatory site. This interaction promotes the ubiquitination of misfolded proteins, presumably through a fourth substrate binding site. As a result, Roq1 alters Ubr1 substrate specificity in a coordinated manner (Figure 7E).

Roq1 induces complex changes in Ubr1, but its architecture is simple. Its two key features, the N-terminal R22 and the YYFV sequence, are so-called short linear motifs (SLiMs). Generally, SLiMs provide monopartite interfaces for protein-protein interactions, even though they are too short to form stable three-dimensional structures. Through their interactions, they act as functional modules and serve as ligands, proteolytic cleavage sites, targeting signals, degrons and sites of posttranslational modification (Davey et al, 2012; Van Roey et al, 2014; Kumar et al, 2024). The two SLiMs of Roq1 are embedded in a completely disordered protein, which makes them maximally accessible. They cooperate in binding Ubr1 as they make physical contacts with distinct sites in Ubr1. The resulting heterobivalent binding mechanism creates avidity and strong Roq1-Ubr1 interaction, even though each SLiM individually is too small to generate high affinity. Such cooperative binding is found in many intrinsically disordered proteins (Holehouse and Kragelund, 2024). In the case of Roq1, cooperative binding is essential not only for a strong overall association with Ubr1 but also to ensure that R22 outcompetes N-degron substrates at the type-1 site and inhibits their degradation.

Remarkably, the Roq1 SLiMs are multifunctional so that diverse biological activities are built into Roq1 by means of only a few amino acid residues. Full-length Roq1 is cleaved by the protease Ynm3 in a relatively well-conserved IL’RSQR sequence (with ’ indicating the cleavage site; Figure S3A). Hence, this sequence is a SLiM that initially serves as a proteolytic cleavage site and, after processing, as a ligand for the Ubr1 type-1 site. The hydrophobic motif has the YYFV sequence at its core, although flanking residues may contribute to Ubr1 binding. Reducing the hydrophobicity of the motif stabilized Roq1, indicating that the motif acts as both a Ubr1 ligand and a degron. Roq1 has a half-life of only about five minutes and its fast turnover may be important for cells to rapidly adjust Roq1 abundance to proteotoxic stress levels (Szoradi et al, 2018).

The region between R22 and the YYFV motif appears to serve solely as a flexible linker. It can be shortened to 15 residues and replaced with generic linkers without disturbing Roq1 function in vitro. Furthermore, it remains possible that an even shorter linker could still maintain function. Flexible linkers can span substantial distances. Assuming a Cα-Cα distance in the peptide backbone of 3.8 Å (Chakraborty et al, 2013), a maximally extended 15-residue linker would, theoretically, be more than 5 nm long. This corresponds to the diameter of a medium-sized globular protein such as actin. Hence, the linker between the Roq1 SLiMs is suited to bridge distant binding sites on the surface of Ubr1.

How could the Roq1 hydrophobic motif enhance the ubiquitination of misfolded proteins? Roq1 does not increase the affinity of Ubr1 for Rad6 or misfolded Pho8*. Instead, it stimulates ubiquitin chain initiation on Pho8* and may additionally stimulate chain elongation. This effect could be explained in several ways. The hydrophobic motif could release autoinhibition of the putative fourth substrate binding site of Ubr1. This mechanism would be similar to how peptide binding to the Ubr1 type-1 and -2 sites releases autoinhibition of the binding site for proteins such as Cup9 (Du et al, 2002). Alternatively or additionally, Roq1 could reshape Ubr1 to optimally position ubiquitin-loaded Rad6 and misfolded substrates for efficient ubiquitin transfer, and Roq1 could increase the flexibility of Ubr1 so that more substrate lysines are accessible for ubiquitination. Resolving these issues will require structural studies of Roq1-Ubr1-substrate complexes. Such studies will also reveal how the hydrophobic motif binds Ubr1 and how Ubr1 binds misfolded proteins. We attempted to determine the structure of the Roq1-Ubr1 complex by cryo-electron microscopy but obtained only low-resolution data. Furthermore, structure prediction for the Roq1-Ubr1 complex by AlphaFold 3 (Abramson et al, 2024) did not yield a binding site for the hydrophobic motif that we could confirm experimentally (unpublished results). However, the interface between the Roq1 hydrophobic motif and Ubr1 is unlikely to involve the Ubr1 type-2 site because disabling this site through point mutations does not disrupt SHRED (Szoradi et al, 2018).

Another challenge is to translate insights from the in vitro studies into an understanding of Ubr1 function in vivo. Ubr1 ubiquitinates the lysine-free Roq1(22-104) in vitro. This modification is dispensable for Roq1 function because Roq1(22-60) is not ubiquitinated and still regulates Ubr1. Nonetheless, our data show that the Rad6-Ubr1 complex can ubiquitinate serines or threonines. Whether this capacity is relevant in vivo needs to be tested. Furthermore, the outcome of Ubr1 reprograming in vivo is likely more complex than can be recapitulated easily in reconstitution experiments. We show that Roq1 activates the ubiquitination of both misfolded proteins and Cup9 in vitro. In contrast, Roq1 appears to suppress Cup9 degradation in vivo (Szoradi et al, 2018). In cells, where various Ubr1 substrates are present simultaneously, Roq1 may stimulate the ubiquitination of certain substrates more strongly than that of others. In effect, this could steer Ubr1 away from proteins that it would readily ubiquitinate if it were not engaged by preferred clients.

Our findings uncover a new mode of ubiquitin ligase regulation, which is defined by cooperating multifunctional motifs in a disordered protein. Elements of this mode have been found in other contexts. Similar to Roq1, the N-termini of human SMAC and YPEL5 block N-degron binding sites of certain ubiquitin ligases as pseudosubstrates, which has been termed degron mimicry (Mace and Day, 2023; Gottemukkala et al, 2024). Importantly, SMAC and YPEL5, through folded domains, make multiple contacts with their partner ubiquitin ligases, and the resulting avidity likely helps them to outcompete the binding of genuine N-degron substrates. The active N-terminus of SMAC is generated by proteolytic processing and SMAC thus also contains a dual-function sequence that is first a proteolytic cleavage site and then an N-degron mimic (Saita et al, 2017). Finally, the proposed action of the SARS-CoV-2 protein ORF10 on the Cullin-RING ubiquitin ligase CUL2 bears intriguing resemblance to the reprograming of Ubr1 by Roq1. ORF10 consists of 38 residues and binds to ZYG11B, a substrate adaptor of CUL2, by means of an N-terminal sequence that mimics a glycine-based N-degron. As a result, the degradation of a glycine/N-degron reporter is inhibited, but the degradation of the intraflagellar transport protein IFT46 is enhanced (Wang et al, 2022; Zhu et al, in press). Hence, there are fascinating similarities between SMAC, YPEL5, ORF10 and Roq1. Intrinsically disordered proteins are often poorly conserved at the sequence level (Holehouse and Kragelund, 2024). In addition, the Roq1 SLiMs consist of only a handful of amino acid residues each. Therefore, it is unsurprising that no recognizable sequence homologs of Roq1 exist in humans. However, the compact elements needed for a Roq1-like regulatory mechanism may be easy to evolve. In fact, intrinsically disordered proteins often acquire SLiMs de novo (Davey et al, 2015). It therefore remains possible that proteins unrelated to Roq1 at the sequence levels but similar in overall construction regulate human Ubr1 homologs.

Ubiquitin ligases are involved in many diseases (Cruz Walma et al, 2022). Artificially controlling or re-directing their activities through inhibitors, molecular glues and proteolysis-targeting chimeras are promising strategies to block or eliminate disease-causing proteins (Cowan and Ciulli, 2022; Tsai et al, 2024; Wang et al, 2024). The regulatory principle embodied by Roq1 requires only a small number of building blocks to achieve far-reaching reprograming of ubiquitin ligases. Therefore, it may be possible to engineer novel Roq1-inspired regulators, potentially also for therapeutic application.

## Methods and Methods

### Plasmids

Plasmids were constructed by Gibson assembly and site-directed mutagenesis and are listed in Table S1. To create pCA528-Roq1(22-104), Roq1(22-104) from pRS316-P_ROQ1_-Roq1 (Szoradi et al, 2018) was inserted into pET24a-His_6_-SUMO (pCA528; Andreasson et al, 2008). Additional insertion of HA or ALFA tags yielded pCA528-Roq1(22-104)-HA and -ALFA. pCA528-Roq1(22-60)-HA was created by replacing Roq1(61-104) of pCA528-Roq1(22-104) with a sequence encoding an HA tag and a stop codon. To generate p416-P_GPD_-Ub-Roq1(22-60), a stop codon was introduced into pRS416-P_GPD_-Ub-Roq1(22-104)-HA(74). To generate pRS316-P_ROQ1_-Ub-Roq1(22-104)-HA(74), an HA tag was inserted into pRS316-P_ROQ1_-Ub-Roq1(22-104) (Szoradi et al, 2018). To create pCA528-Pho8*-MBP, Pho8* from pRS305-P_ADH_-Rtn1-Pho8*-FLAG-GFP (Szoradi et al, 2018) was inserted into pCA528, generating pCA528-Pho8*. Subsequently, MBP from pMal-c2E-Hsp42-FLAG (Ungelenk et al, 2016) was modified with an upstream sequence encoding a GGSGGGSGG linker and inserted into pCA528-Pho8*. pCA528-Pho8-MBP was created by reverting the F352S mutation in pCA528-Pho8*-MBP. pCA528-R-GFP and pCA528-F-GFP were generated by inserting codons for arginine or phenylalanine and a sequence encoding a SKGEELFYGV linker into pCA528-GFP (Schmidt et al, 2009). To generate pCA528-Cup9-Strep, Cup9 from yeast genomic DNA was modified with a Strep tag II and inserted into pCA528. To generate pCA528-Mgt1-MBP, Mgt1 from yeast genomic DNA was inserted into pCA528-Pho8*-MBP, replacing Pho8*. To generate YEplac181-P_ADH_-Chk1-ALFA-FLAG, Chk1 from yeast genomic DNA was modified with an ALFA-FLAG tag and inserted into YEplac181-P_ADH_-FLAG-Ubr1 (pFLAGUBR1SBX; Xia et al., 2008), replacing FLAG-Ubr1.

### Protein purification

Ubr1 was purified essentially as described (Du et al, 2002). Protease-deficient yeast expressing FLAG-Ubr1 (SSY2908) were grown to OD_600_ = 2 at 30°C in 12 L YPD medium, washed with cold PBS, resuspended in 6 x 10 ml lysis buffer (50 mM HEPES pH 7.5, 0.2 M NaCl, 10% glycerol, 0.5% NP-40, 1 mM PMSF, Roche complete protease inhibitors) and disrupted by 10 passages through a Microfluidics M-110L microfluidizer at 1200 bar. Lysates were cleared at 11,200 g for 30 min and the supernatant was incubated with 0.9 ml anti-FLAG agarose beads (Thermo Fisher Scientific) at 4°C for 2 h. Beads were washed with 20 ml buffer A (50 mM HEPES pH 7.5, 1 M NaCl, 10 % glycerol, 0.5% NP-40) and 20 ml buffer B (50 mM HEPES pH 7.5, 0.2 M NaCl, 10% glycerol), and FLAG-Ubr1 was eluted with 5 x 500 µl buffer B containing 1 mg/ml FLAG peptide (Thermo Fisher Scientific). Protein-containing eluates were pooled and concentrated with an Amicon Ultra centrifugal filter with a 100 kDa molecular weight cut-off.

Chk1 was purified from protease-deficient yeast (PWY0005) harboring YEPlac181-P_ADH_-Chk1-ALFA-FLAG. Cells were grown to OD_600_ = 2 at 30°C in SCD medium without leucine, and Chk1-ALFA-FLAG was isolated as described for FLAG-Ubr1.

Roq1 variants were purified from *E. coli* BL21(DE3) harboring pCA528-based expression plasmids for Roq1 with an N-terminal His_6_-SUMO tag. Cells were grown to OD_600_ = 0.7 at 37°C in 6 L 2xYT medium with 25 µg/ml kanamycin and 34 µg/ml chloramphenicol, and Roq1 expression was induced with 1 mM IPTG (ZellBio) at 37°C for 3 h. Cells were harvested at 1000 g at 4°C for 15 min, washed with cold water and resuspended in 60 ml lysis buffer (50 mM Bis-Tris pH 6.0, 500 mM NaCl, 20 mM imidazole, 5 mM MgCl_2_, 2 mM 2-mercaptoethanol [2-ME], 1 mM PMSF, Roche protease inhibitors without EDTA). A dollop of DNase (Merck) was added and the cell suspension was stirred at 4°C for 20 min. Cells were disrupted by 6 passages through an M-110L microfluidizer and the lysate was cleared at 11,200 g at 4°C for 30 min. For immobilized metal affinity chromatography (IMAC), the supernatant was incubated with 1.5 g Ni-IDA beads (Macherey-Nagel) for 60 min. The beads were applied to a gravity flow column, washed with lysis buffer, wash buffer (50 mM Bis-Tris pH 6.0, 50 mM imidazole, 500 mM NaCl, 5 mM MgCl_2_, 2 mM 2-ME), wash buffer with 750 mM NaCl and wash buffer with 5 mM ATP. Bound protein was eluted with elution buffer (50 mM Bis-Tris pH 6.0, 150 mM NaCl, 250 mM imidazole, 2 mM 2-ME) and protein-containing fractions were pooled. To remove the N-terminal His_6_-SUMO tag, the eluate was mixed with 25 µl His-tagged SUMO protease Ulp1 (purified as described below) and dialyzed with a 3 kDa molecular weight cut-off against dialysis buffer (50 mM Bis-Tris pH 6.0, 150 mM NaCl, 2 mM 2-ME) at 4°C overnight. To remove cleaved His_6_-SUMO, uncleaved His_6_-SUMO-Roq1 and Ulp1-His_6_, Ni-IDA beads were added for 30 min and collected with a gravity flow column. The flow-through was concentrated and applied to a Hiload 16/600 Superdex S30 prep grade size exclusion column (Cytiva) equilibrated with 50 mM HEPES pH 7.5, 150 mM NaCl. Protein-containing fractions were pooled and 10% (v/v) glycerol were added.

Roq1 for photo-crosslinking was expressed with the pEVOL/pET system (Young et al, 2010). *E. coli* BL21(DE3) harboring pEVOL-pBpF and pCA528-Roq1(22-104)-ALFA with Y55amber or Y56amber mutations were grown in 2 L 2xYT medium containing antibiotics, 50 mM phosphate buffer pH 7.2 and 1 mM p-benzoyl-L-phenylalanine (Bachem). L-arabinose (Merck) was added to 0.25% (w/v) when cultures reached OD_600_ = 0.5 and again at OD_600_ = 0.8. Roq1 expression was induced with IPTG at 37°C for 5 h, cells were lysed, His_6_-SUMO-Roq1-ALFA was purified by IMAC and cleaved with Ulp1 as described above. To immobilize Roq1, 16 ml dialysate were incubated with 250 µl magnetic ALFA Selector CE beads (50% slurry, NanoTag Biotechnologies) at 4°C for 2 h. Beads were washed with 3 x 500 µl wash buffer (50 mM HEPES, pH 7.5, 0.15 M NaCl, 10 mM MgCl_2_) and used for photo-crosslinking.

To purify Rad6, *E. coli* BL21(DE3) harboring pCA528-Rad6 were grown in 2xYT medium with antibiotics, treated with IPTG at 37°C for 4 h and lysed as above in 50 mM NaH_2_PO_4_ pH 8.0, 300 mM NaCl, 5 mM MgCl_2_, 2 mM 2-ME, 10 mM imidazole and Roche protease inhibitors without EDTA. After removal of the His_6_-SUMO tag, Rad6 was further purified on a Hiload 16/60 Superdex S75 prep grade size exclusion column (GE HealthCare) equilibrated with 25 mM HEPES pH 7.5, 25 mM KCl, 5 mM MgCl_2_, 1 mM DTT. Protein-containing fractions were pooled and 10% (v/v) glycerol were added.

R-GFP and F-GFP were purified from *E. coli* BL21(DE3) harboring pCA528-R-GFP or pCA528-F-GFP. The same protocol was used as for the purification of Rad6 except that a Hiload 16/60 Superdex S200 prep grade column (GE HealthCare) was used.

To purify Pho8 and Pho8*, *E. coli* BL21(DE3) harboring pCA528-Pho8-MBP or pCA528-Pho8*-MBP were grown in LB medium containing 25 µg/ml kanamycin and 34 µg/ml chloramphenicol, treated with IPTG at 30°C for 3 h and lysed as above in 50 mM HEPES pH 7.5, 500 mM NaCl, 5 mM MgCl_2_, 10% glycerol, 10 mM imidazole, 0.4% CHAPS, 1 mM PMSF and Roche protease inhibitors without EDTA. Lysates were cleared and the supernatant was applied to a HisTrap FF crude affinity chromatography column (GE HealthCare) equilibrated with IMAC buffer (50 mM HEPES pH 7.5, 500 mM NaCl, 5 mM MgCl_2_, 10% glycerol). The column was washed with 10 volumes IMAC buffer containing 10 mM imidazole and 10 mM ATP, and protein was eluted with IMAC buffer containing 250 mM imidazole. The eluate was mixed with Ulp1 and dialyzed overnight against dialysis buffer (50 mM HEPES pH 7.5, 150 mM NaCl, 10 mM MgCl_2_, 2 mM 2-ME). The dialysate was further batch-purified with amylose resin (NEB) equilibrated with dialysis buffer, bound protein was eluted with 10 mM maltose in dialysis buffer, the eluate was concentrated and loaded onto a Hiload 16/60 Superdex S200 size exclusion prep grade column equilibrated with 50 mM HEPES pH 7.5, 150 mM NaCl, 10 mM MgCl_2_. Protein-containing fractions were pooled and 10% (v/v) glycerol were added.

To purify Cup9, *E. coli* BL21(DE3) harboring pCA528-Cup9-Strep were grown in LB medium with antibiotics, treated with IPTG at 37°C for 3 h and lysed in 20 mM HEPES pH 7.0, 500 mM NaCl, 10 mM imidazole, 5 mM MgCl_2_, 2 mM 2-ME, 1 mM PMSF and Roche protease inhibitors without EDTA. IMAC purification and Ulp1 digest were done as described for Roq1. The dialysate was loaded onto a StrepTrap HP column (Cytiva) equilibrated with Strep buffer (20 mM HEPES pH 7.0, 500 mM NaCl). The column was washed with 10 volumes Strep buffer, Cup9-Strep was eluted with 2.5 mM D-desthiobiotin in Strep buffer with 2 mM 2-ME and applied to a Hiload 16/60 Superdex S200 size exclusion prep grade column equilibrated with 20 mM HEPES pH 7.5, 500 mM NaCl, 1 mM DTT. Protein-containing fractions were pooled and 10% (v/v) glycerol were added.

To purify Mgt1, *E. coli* BL21(DE3) harboring pCA528-Mgt1-MBP were grown in LB medium with antibiotics, treated with IPTG at 30°C for 3 h and lysed as above. After IMAC, protein-containing fractions were pooled and batch-purified with amylose resin (NEB) equilibrated with 50 mM HEPES pH 7.5, 500 mM NaCl. The column was washed and protein was eluted with 10 mM maltose, digested with Ulp1, dialyzed, concentrated and applied to a Hiload 16/60 Superdex S75 prep grade size exclusion column (GE HealthCare) equilibrated with 50 mM HEPES pH 7.5, 150 mM NaCl, 10 mM MgCl_2_. Protein-containing fractions were pooled and 10% (v/v) glycerol were added.

To purify Ulp1 protease, *E. coli* BL21(DE3) harboring pET24a-Ulp1-His_6_ were grown at 30°C in 2xYT medium containing 100 µg/ml ampicillin and 34 µg/ml chloramphenicol, and Ulp1 expression was induced with 0.5 mM ITPG at 20°C overnight. Cells were lysed in 20 mM Tris pH 7.9, 100 mM KCl, 0.6% Brij-58, 5 mM MgCl_2_, 2 mM 2-ME, 1 mM PMSF and Roche protease inhibitors without EDTA. Ulp1-His_6_ was immobilized on Ni-IDA beads, washed with 300 ml wash buffer A (20 mM Tris pH 7.9, 2 M urea, 100 mM KCl, 0.1% Brij-58, 2 mM 2-ME), 300 ml buffer B (20 mM Tris pH 7.9, 1 M KCl, 0.1% Brij-58, 2 mM 2-ME) and 300 ml buffer C (20 mM Tris pH 7.9, 100 mM KCl, 2 mM 2-ME) and eluted with lysis buffer containing 250 mM imidazole. Protein-containing fractions were pooled and dialyzed overnight against dialysis buffer (40 mM HEPES pH 7.9, 150 mM KCl, 10% glycerol, 2 mM 2-ME). Aggregates were removed by centrifugation at 4000 rpm at 4°C for 15 min, protein was concentrated to 8 mg/ml and 50% (v/v) glycerol were added.

To purify untagged ubiquitin, *E. coli* BL21(DE3) harboring pET3a-ubiquitin were grown in LB medium with 100 µg/ml ampicillin and 34 µg/ml chloramphenicol, treated with 1 mM IPTG at 37°C for 4 h and lysed in 50 mM Tris-HCl pH 7.6, 0.02% NP-40, 10 mM MgCl_2_, 1 mM DTT, 1 mM PMSF and Roche complete protease inhibitors. Proteins were precipitated with cold 70% perchloric acid and separated from soluble ubiquitin by centrifugation. The supernatant was dialyzed against 50 mM ammonium acetate pH 4.5. Ubiquitin was further purified by cation exchange on a Resource S column (Cytiva) equilibrated with dialysis buffer and eluted with a linear gradient of 0 - 0.5 M NaCl in 50 mM ammonium acetate pH 4.5 over 20 column volumes. Protein-containing fractions were pooled, concentrated and applied to a Hiload 16/60 Superdex S75 prep grade size exclusion column (GE HealthCare) equilibrated with 50 mM Tris-HCl pH 7.6, 150 mM NaCl, 1 mM DTT.

### Chemical unfolding of Luciferase

To obtain unfolded but soluble Luciferase^U^, recombinant firefly Luciferase (Promega) was diluted four-fold in denaturation buffer (30 mM Tris pH 7.4, 50 mM KCl, 5 mM MgCl_2_, 7.5 M guanidinium chloride, 10 mM DTT) and incubated on ice for 30 min. Luciferase was then diluted 100-fold in stabilization buffer (30 mM Tris pH 7.4, 50 mM KCl, 5 mM MgCl_2_, 0.2 M trehalose, 1 mM DTT) according to Singer and Lindquist, 1998. To eliminate aggregates, samples were centrifuged at 13,200 g at 4°C for 30 min and the supernatant was used for in vitro ubiquitination assays. To obtain folded Luciferase^N^, recombinant firefly Luciferase was treated as above except that guanidinium chloride was omitted.

### In vitro ubiquitination assays

Ubiquitination assays were done in 10 µl containing 0.1 µM Ube1 (R&D Systems), 4 µM Rad6, 0.25 µM Ubr1, 80 µM wild-type or lysine-free ubiquitin (MoBiTec), and 0.2 µM substrate protein in reaction buffer (50 mM HEPES pH 7.5, 0.15 M NaCl, 10 mM MgCl_2_, 5 mM ATP). Roq1 variants were used at 2.5 µM unless stated otherwise. The RA dipeptide (peptides & elephants) was used at a final concentration of 1 mM. Reactions were incubated at 30°C for the times indicated and stopped with SDS-PAGE sample buffer containing DTT. To hydrolyze ester bonds between ubiquitin and substrate proteins, ubiquitination reactions were set up as above and incubated at 30°C for 30 min. SDS-PAGE sample buffer was added and samples were incubated at 65°C for 5 min. NaOH was added to 0.25 mol/L, samples were incubated at room temperature for 10 min, NaOH was neutralized with 0.25 mol/L HCl and samples were used for western blotting. Control samples were treated as above except that NaOH and HCl were replaced with water.

### Western blotting

Cells were collected by centrifugation, washed with water, resuspended in lysis buffer (50 mM HEPES pH 7.5, 0.5 mM EDTA, 1 mM PMSF, Roche complete protease inhibitors), and disrupted by bead beating with a FastPrep 24 (MP Biomedicals). Proteins were solubilized by addition of 1.5% (w/v) SDS and incubation at 65°C for 5 min. Lysates were cleared at 16,000 g at 4°C for 2 min and protein concentrations were determined with a BCA kit (Pierce). Proteins were separated by Tris-glycine or Tris-tricine SDS-PAGE and transferred onto nitrocellulose membranes by wet blotting. Membranes were probed with primary and HRP-coupled secondary antibodies, HRP conjugates or fluorescently labeled nanobodies (Table S2). For chemiluminescence detection, membranes were incubated with SuperSignal West Pico Plus substrate (Thermo Fisher Scientific) and analyzed with an Amersham Imager 600 (GE HealthCare). For fluorescence detection, membranes were analyzed with an Odyssey CLx imaging system (LI-COR).

### Pho8* solubility assay

To determine Pho8* solubility, duplicate 10 µl ubiquitination assays including Roq1 were set up as described above. After 0 or 90 min, samples were saved as input or centrifuged at 18,000 g at 4°C for 30 min. Supernatants were collected and pellets were resuspended in 10 µl reaction buffer (50 mM HEPES, pH 7.5, 0.15 M NaCl, 10 mM MgCl_2_). Equal volumes of input, supernatant and pellet samples were analyzed by western blotting.

### Biolayer Interferometry

The Roq1-Ubr1 interaction was analyzed with the OctetRed96e system (Sartorius). Roq1(22-104) variants were biotinylated at C103, the single cysteine in Roq1, by incubation with a 1.2-fold molar excess of thiol-reactive EZ-Link^TM^ HPDP-Biotin (Thermo Fisher Scientific) at room temperature for 90 min. Unbound biotin was removed with a Zeba spin desalting column (Thermo Fisher Scientific) and biotinylated Roq1 was immobilized on streptavidin biosensors (Sartorius) that had been hydrated in assay buffer (50 mM HEPES, 150 mM NaCl, 10 mM MgCl_2_, 0.05 % Tween-20, 0.02 % BSA) for 10 min. Binding assays were performed in black 96-well plates (Greiner) in 200 µl at 1000 rpm. Assays consisted of (1) a washing step in assay buffer for 1 min, (2) a loading step with 1.25 µg/ml wild-type Roq1 or mutant variants for 10 min, (3) a washing step for 1 min, (4) a baseline step for data normalization for 1 min, and (5) an association step with 50 nM Ubr1 (Figures 4C and 5A) or 100 nM Ubr1 (Figure 2D) for 5 min. Rad6-Ubr1 interaction was analyzed with Rad6 biotinylated with a 1.2-fold molar excess of the amino-reactive EZ-Link^TM^ NHS-PEG4-Biotin (Thermo Fisher Scientific) at room temperature for 30 min. The set-up of the assay was the same as above except that 0.5 µg/ml Rad6 were used for loading. To test the effect of Roq1 on Rad6-Ubr1 association, 100 nM Ubr1 and 200 nM or 1 µM Roq1 were used. The Roq1-Ubr1 and Rad6-Ubr1 interactions were essentially irreversible so that only the association could be analyzed.

### Bioinformatic analyses

The structure prediction for full-length Roq1 was retrieved from the AlphaFold Protein Structure Database (Varadi et al, 2024). The disorder prediction was obtained by averaging estimates for the disorder tendency across the full-length Roq1 sequence from 12 prediction algorithms, namely AIUPred, DisEMBL, DISpro, Espritz, IsUnstruct, IUPred3, Metapredict, MFDp, NetsurfP 3.0, PONDR VSL2, PrDOS and RONN (Kurgan, 2022). To identify Roq1 sequence homologs, a protein-protein BLAST search excluding *S. cerevisiae* was carried out via the NCBI home page. An alignment of the retrieved sequences from *Saccharomyces* and *Kazachstania* species was done with CLUSTALW.

### Yeast strains

Strains used in this study were derived from W303 mating type a and are listed in Table S3. To generate strain SSY2959, the P_ADH_-FLAG-Ubr1 cassette was amplified from plasmid pRS415-P_ADH_-FLAG-Ubr1 and integrated into the *leu2* locus of SSY792. To generate SSY3543, 3544, 3545 and 3551, the expression cassettes of pRS306N-derived plasmids were linearized by PCR and integrated into the *ura3* locus of SSY835.

### Roq1 mutagenesis screen

Roq1-HA(74) was amplified from pRS416-P_GPD_-Roq1-HA(74) (Szoradi et al, 2018) by error-prone PCR. Reactions consisted of 1 ng/µl plasmid DNA, 0.5 µM primer GPDprom-165, 0.5 µM primer CYCterm-66, 200 µM dATP, 200 µM dGTP, 1 mM dTTP, 1 mM dCTP, 1.5 mM MgCl_2_, 0.25 mM MnCl_2_ and 0.04 U/µl Taq polymerase (Sigma, D1806) in PCR buffer without MgCl_2_ (Sigma, D4545). Sequencing of PCR products showed that these conditions resulted in about 0.4 mutations per 600-bp amplicon. PCR products were then introduced into yeast by gap repair cloning. Roq1 was excised from pRS416-P_GPD_-Roq1-HA(74) by restriction digest with SpeI, XhoI and BseRI, and the plasmid backbone was used, together with the products of the error-prone PCR, for co-transformation of the SSY835 reporter strain. Transformants were plated onto SCD-URA plates and about 20,000 colonies were visually screened for impaired degradation of the Rtn1-Pho8*-GFP reporter as judged by retention of GFP fluorescence. Candidates were validated with the flow cytometry-based SHRED assay described below. To exclude mutations that destabilized the Roq1 protein, remaining candidates were tested for Roq1 expression by western blotting. Plasmids from candidates with detectable Roq1 expression were sequenced, which revealed 14 unique point mutations in the Roq1 coding sequence. To confirm that the defects in reporter degradation were caused by mutations in plasmid-encoded Roq1 rather than chromosomal mutations, the point mutations were introduced into pRS416-P_GPD_-Roq1(22-104)-HA(74), the resulting plasmids were introduced into SSY835 and SHRED activity was measured by flow cytometry.

### SHRED assay

Yeast expressing the SHRED reporter Rtn1-Pho8*-GFP and cytosolic BFP were grown to mid-log phase in 1 ml SCD medium in 96 deep-well plates at 30°C. Cultures were diluted to OD_600_ = 0.05 and were left untreated or treated with 2 µg/ml tunicamycin (Merck) for 5 h. Fluorescence was measured with a FACS Canto flow cytometer equipped with a high-throughput sampler (BD Biosciences). To determine reporter levels, GFP fluorescence was first corrected for autofluorescence by subtracting the fluorescence of identically treated control cells not expressing GFP. Corrected GFP fluorescence was divided by BFP fluorescence to normalize for cellular translation capacity. To determine the effect of stress treatment, GFP/BFP ratios of treated cells were divided by those of corresponding untreated cells. The resulting reporter levels were expressed in per cent of those at t = 0. Roq1 variants lacking the first 21 amino acid residues were expressed as N-terminal ubiquitin fusions, which, after processing by endogenous ubiquitin proteases, yielded Roq1 cleavage fragments starting with R22.

### Immunoprecipitation

Twenty ODs of yeast cells in mid-log phase were lysed by bead beating in IP buffer (25 mM HEPES pH 7.5, 100 mM NaCl, 1% Triton X-100, 0.5 mM EDTA, 1 mM PMSF, Roche complete protease inhibitors). Lysates were cleared at 12,000 g at 4°C for 5 min. 3% of the lysate were kept to analyze Roq1 in the input, 15% were kept to analyze Ubr1 in the input and 50% were used to precipitate Roq1 with 30 µl anti-HA-agarose beads (Sigma) at 4°C for 30 min. Beads were washed three times with cold lysis buffer and bound protein was eluted with SDS-PAGE sample buffer at 65°C for 5 min. For the analysis of Ubr1 in the input sample, Ubr1 was first immunoprecipitated with 20 µl anti-FLAG agarose beads (clone M2, Sigma). This step reduced variability between samples, which occurred when Ubr1 levels were analyzed directly by western blotting of whole cell lysates.

### Photo-crosslinking

Roq1(22-104)-ALFA and variants containing p-benzoyl-L-phenylalanine were immobilized on ALFA Selector CE beads (see above) and incubated with 150 µl wash buffer (50 mM HEPES, pH 7.5, 0.15 M NaCl, 10 mM MgCl_2_) containing 170 nM FLAG-Ubr1 at 4°C overnight to allow formation of the Roq1-Ubr1 complex. Beads were washed with wash buffer and Roq1 was eluted with 2 x 25 µl wash buffer containing 800 µM ALFA peptide (NanoTag Biotechnologies). 20 µl eluate were kept as control sample and another 20 µl were used for photo-crosslinking. Samples were irradiated on ice with a UV-LED lamp (Opsytec Dr. Gröbel GmbH) at 365 nm with 15 cycles of 1 s irradiation at 25 W/cm^2^ followed by a 2 s pause. Control and irradiated samples were then resolved on a 7.5/15% step gradient Tris-glycine SDS-PAGE gel and analyzed by western blotting.

### In vitro pulldown

To assess Pho8/Pho8*-Ubr1 interaction, 0.25 µM Pho8-MBP/Pho8*-MBP, 0.25 µM FLAG-Ubr1 and, where indicated, 2.5 µM Roq1(22-60)-HA in 30 µl pulldown buffer (50 mM HEPES pH 7.5, 150 mM NaCl, 10 mM MgCl_2_, 0.66 µg/µl BSA) were incubated at 30°C for 90 min. 12 µl amylose resin (NEB) equilibrated in pulldown buffer were added and samples were incubated with rotation at 4°C for 2 h. Beads were washed for 3 x 5 minutes with 100 µl wash buffer (50 mM HEPES pH 7.5, 150 mM NaCl, 10 mM MgCl_2_), protein was eluted with 2 x 10 µl elution buffer (50 mM HEPES pH 7.5, 150 mM NaCl, 10 mM MgCl_2_, 10 mM maltose) at 4°C for 15 min and eluates were analyzed by western blotting.

## Acknowledgements

We are indebted to Chi-Ting Ho, Ilia Kats, Jörg Malsam, Matthias Mayer, Frauke Melchior, Carola Sparn for reagents, protocols and advice. We thank Claudio Joazeiro, Matthias Mayer, Dimitris Papagiannidis, Anne Schlaitz and all Schookees for comments on the manuscript. This work was supported by grant SCHU 2364/2-1 from the German Research Foundation (DFG) to SS and a fellowship from the Boehringer Ingelheim Fonds to NP. The authors gratefully acknowledge the data storage service SDS@hd supported by the Ministry of Science, Research and the Arts Baden-Württemberg (MWK) and the DFG through grant INST 35/1503-1 FUGG. The authors declare no competing financial interests.

## Author contributions

Conceptualization: Sibylle Kanngießer, Niklas Peters, Sebastian Schuck; Investigation: Sibylle Kanngießer, Oliver Pajonk, Niklas Peters, Rafael Salazar Claros; Resources: Axel Mogk; Supervision: Sebastian Schuck; Writing - original draft: Sibylle Kanngießer, Niklas Peters, Sebastian Schuck; Writing - review and editing: all authors.

---

**Table S1.**
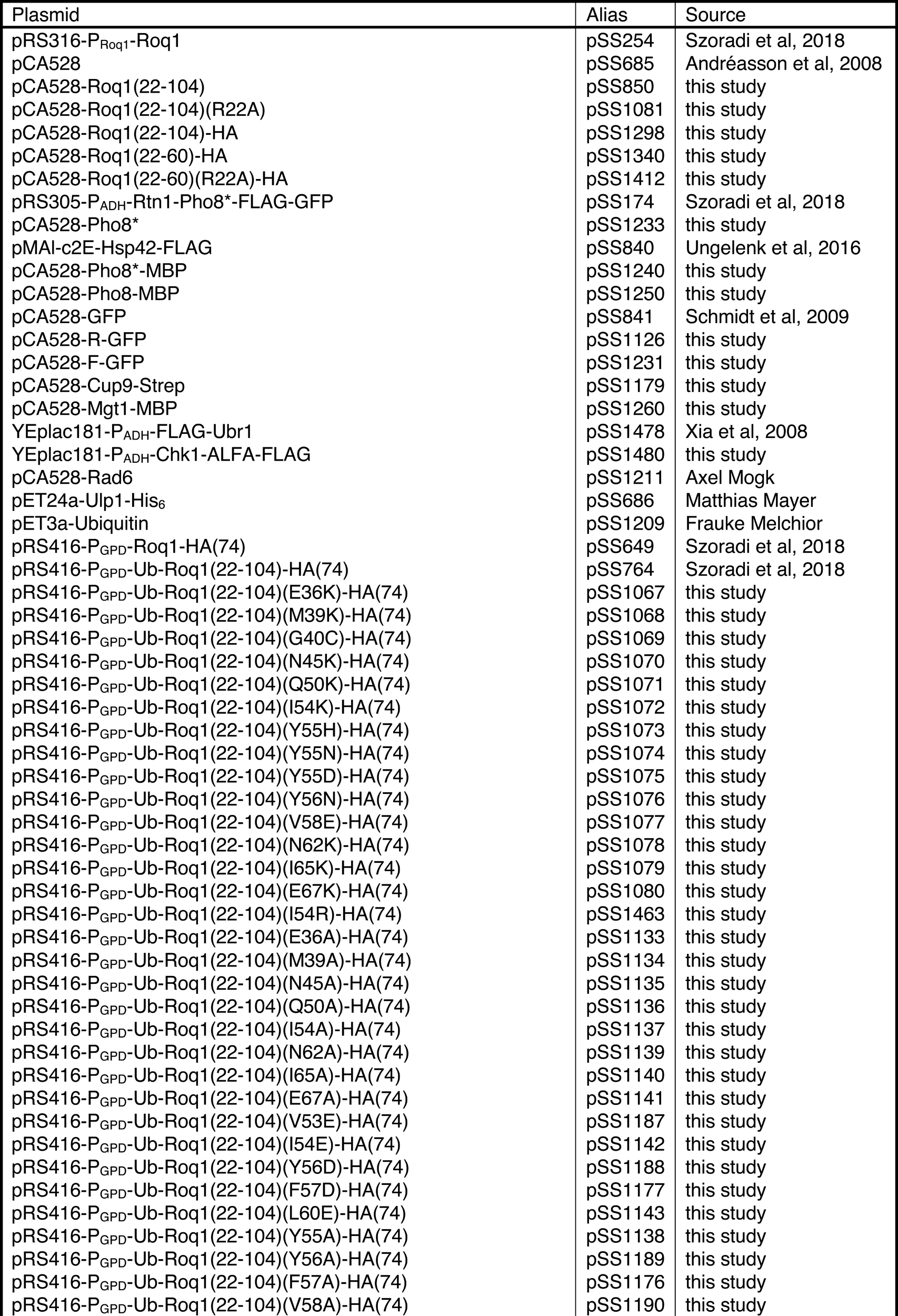
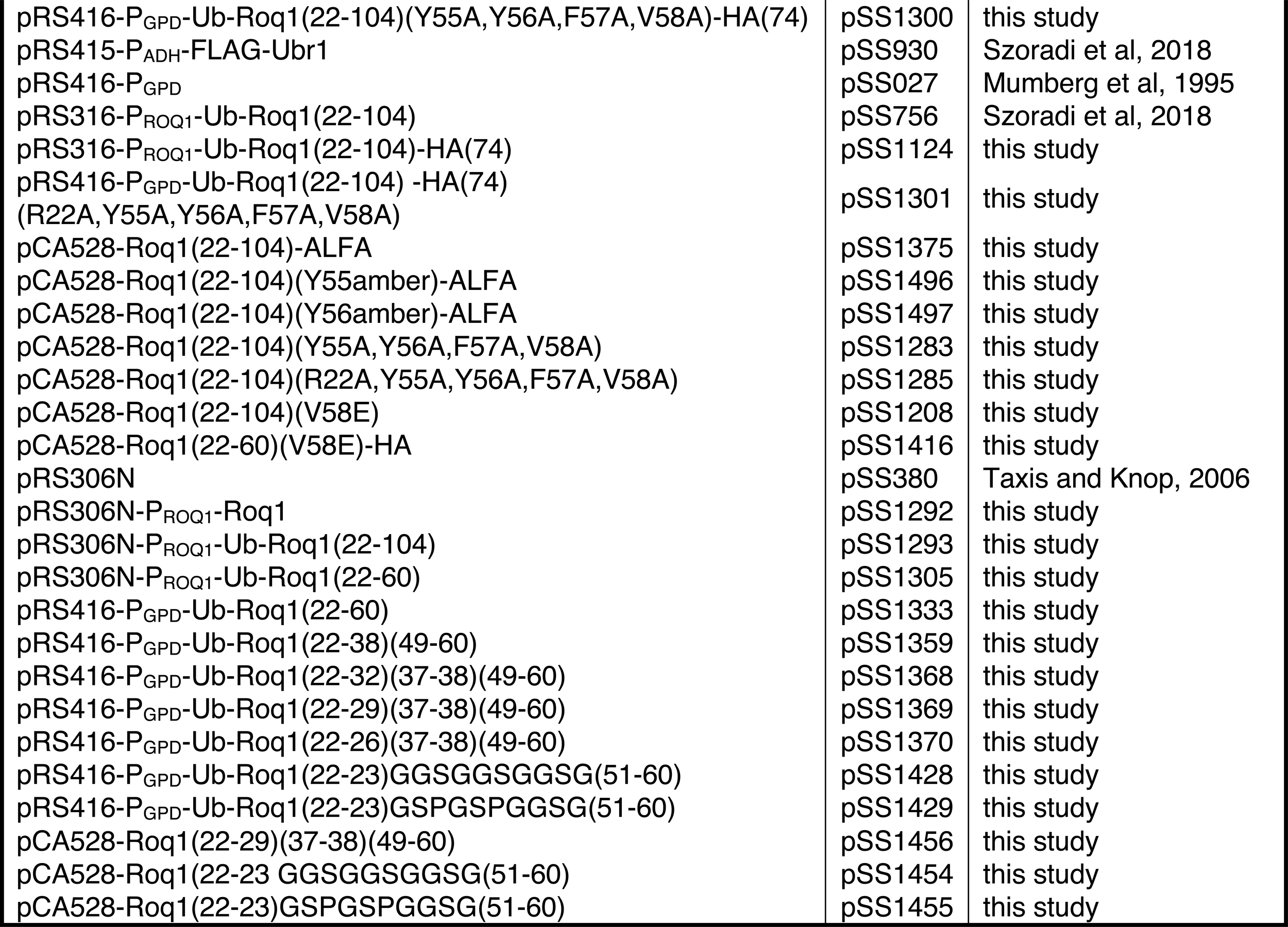
Plasmids used in this study.

**Table S2.** Antibodies and conjugates used in this study. HRP = horseradish peroxidase; cat# = catalog number; RRID = research resource identifier.

| Antibody/conjugate | Source | Identifier (cat#; RRID) |
| --- | --- | --- |
| rat anti-HA (clone 3F10) | Roche | 11867423001; AB_390918 |
| mouse anti-Pho8 (clone 1D3A10) | Abcam | Ab113688; AB_10860792 |
| goat anti-Luciferase | Merck | 3256 |
| mouse anti-GFP (clones 7.1/13.1) | Roche | 11814460001; AB_390913 |
| mouse anti-FLAG (clone M2) | Merck | F1804; AB_262044 |
| mouse anti-MBP (clone MBP-17) | Merck | M6295 |
| Strep-Tactin HRP | Iba Lifesciences | 2-1502-001 |
| donkey anti-rat HRP | Jackson ImmunoResearch | 712-035-153; AB_2340639 |
| goat anti-mouse HRP | Thermo Fisher Scientific | 31432; AB_228302 |
| rabbit anti-goat HRP | Jackson ImmunoResearch | 305-035-045; AB_2339403 |
| anti-ALFA IRDye 800CW | NanoTag Biotechnologies | N1502-Li800-L; AB_3075985 |

**Table S3.**
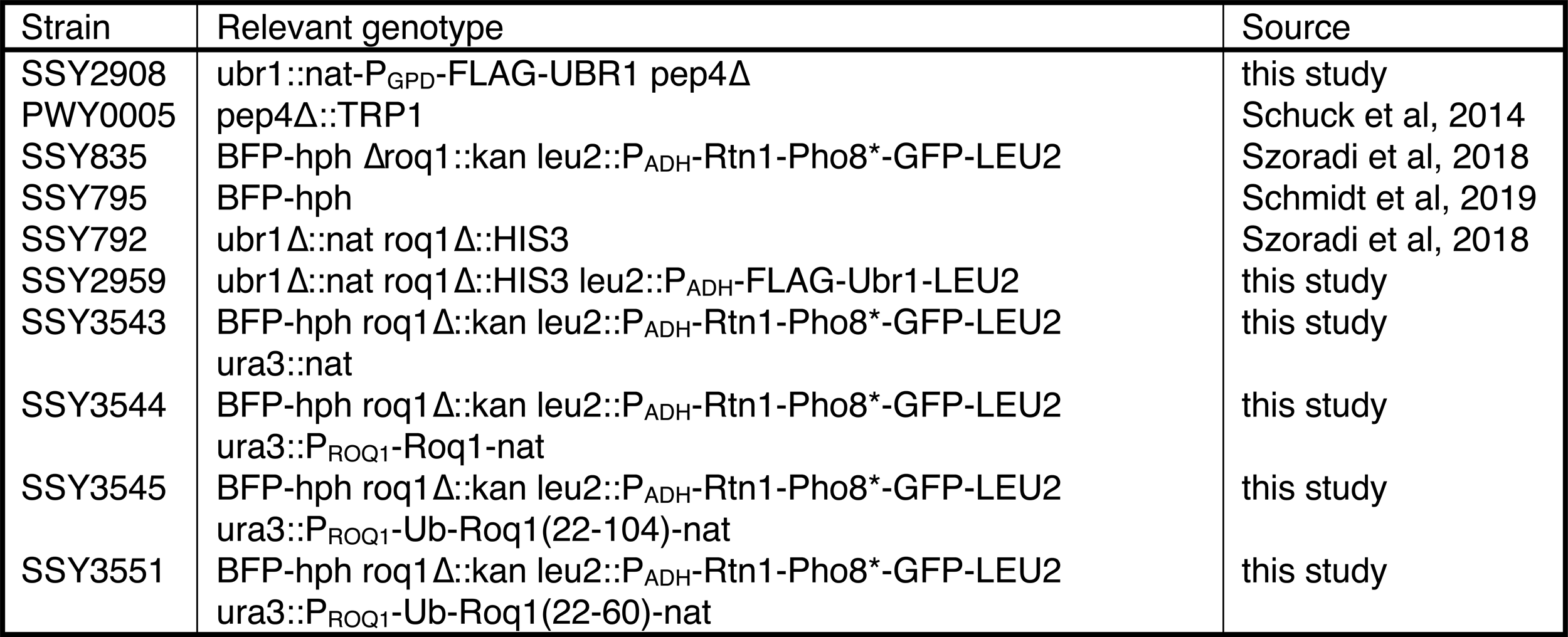
Yeast strains used in this study. Only the chromosomal genotype is given for strains harboring episomal plasmids. BFP-hph = his3::P_GPD_-BFP-hph.

**Figure S1.**
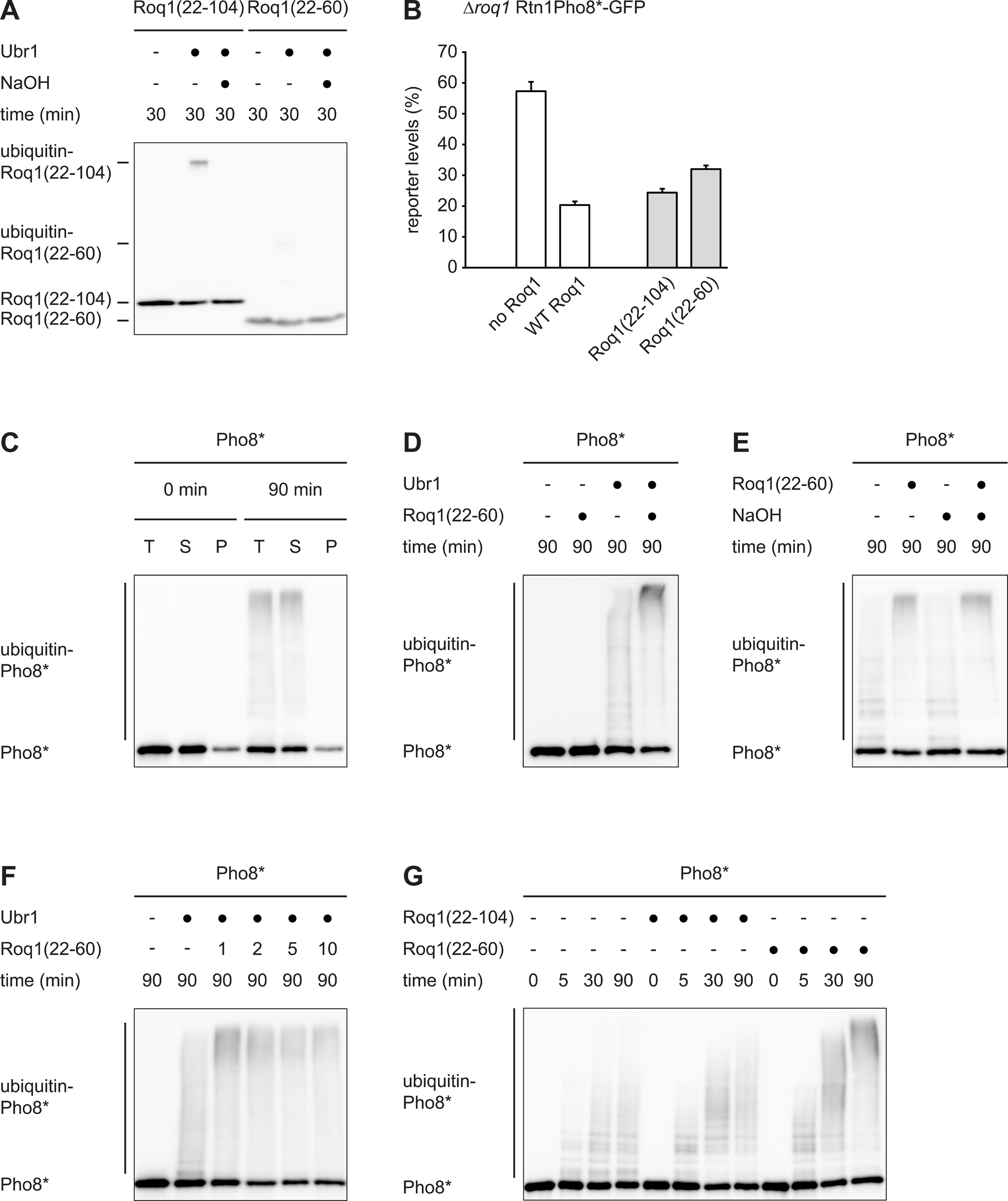
In vitro reconstitution of Ubr1 regulation by Roq1. A Western blot of HA tag from ubiquitination assays with Roq1(22-104)-HA and Roq1(22-60)-HA in the absence of a designated Ubr1 substrate. Ubiquitination reactions were stopped after 30 min with buffer containing dithiothreitol and, where indicated, treated with NaOH to hydrolyze ester bonds between Roq1 and ubiquitin. Ubr1 catalyzes the formation of NaOH-sensitive conjugates of Roq1(22-104) and ubiquitin, which is almost completely lost by shortening Roq1 to Roq1(22-60). B Cellular levels of the SHRED reporter Rtn1-Pho8*-GFP after tunicamycin treatment for 5 h relative to levels in untreated cells, as measured by flow cytometry. The *roq1* mutant cells contained an empty plasmid (no Roq1), a plasmid encoding wild-type Roq1 (WT Roq1), ubiquitin-Roq1(22-104) or ubiquitin-Roq1(22-60). The ubiquitin fusions are processed by cells to yield Roq1(22-104) or Roq1(22-60) starting with R22. Bars are the mean ± SEM; n = 3 biological replicates. C Western blot of Pho8 from solubility assays of Pho8*. Ubiquitination assays including Roq1(22-60) were carried out for 0 or 90 min and soluble and insoluble Pho8* were separated by centrifugation. T = total; S = supernatant; P = pellet. D Western blot of Pho8 from Pho8* ubiquitination assays with and without Ubr1 and Roq1(22-60). No Pho8* ubiquitination occurs in the absence of Ubr1. E Western blot of Pho8 from Pho8* ubiquitination assays with and without Roq1(22-60). Ubiquitination reactions were stopped after 90 min and, where indicated, treated with NaOH to hydrolyze ester bonds between Pho8* and ubiquitin. Ubiquitin-Pho8* conjugates were resistant to alkaline hydrolysis, showing that they consisted of amide rather than oxyester bonds. F Western blot of Pho8 from Pho8* ubiquitination assays with different concentrations of Roq1(22-60). Roq1(22-60) was omitted or used at molar ratios of 1:1, 2:1, 5:1 and 10:1 relative to Ubr1. G Western blot of Pho8 from Pho8* ubiquitination assays without and with Roq1(22-104) or Roq1(22-60) for the times indicated.

**Figure S2.**
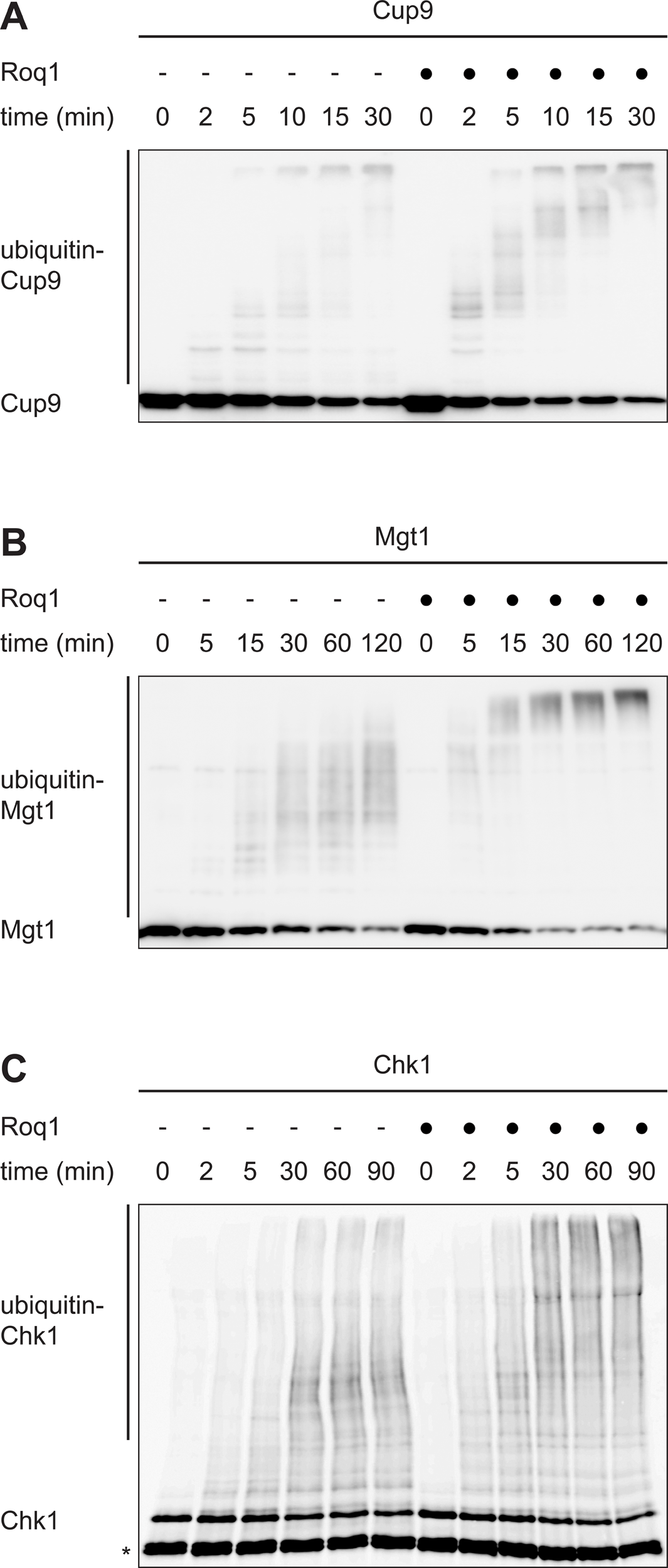
Roq1 stimulates ubiquitination of folded Ubr1 substrate proteins with internal degrons. A Western blot of Strep tag from Cup9-Strep ubiquitination assays without and with Roq1(22-60) for the times indicated. B Western blot of maltose-binding protein (MBP) from Mgt1-MBP ubiquitination assays without and with Roq1(22-60) for the times indicated. C Western blot of FLAG tag from Chk1-ALFA-FLAG ubiquitination assays without and with Roq1(22-60) for the times indicated. The asterisk denotes truncated Chk1 that arose during the expression and purification of Chk1-ALFA-FLAG.

**Figure S3.**
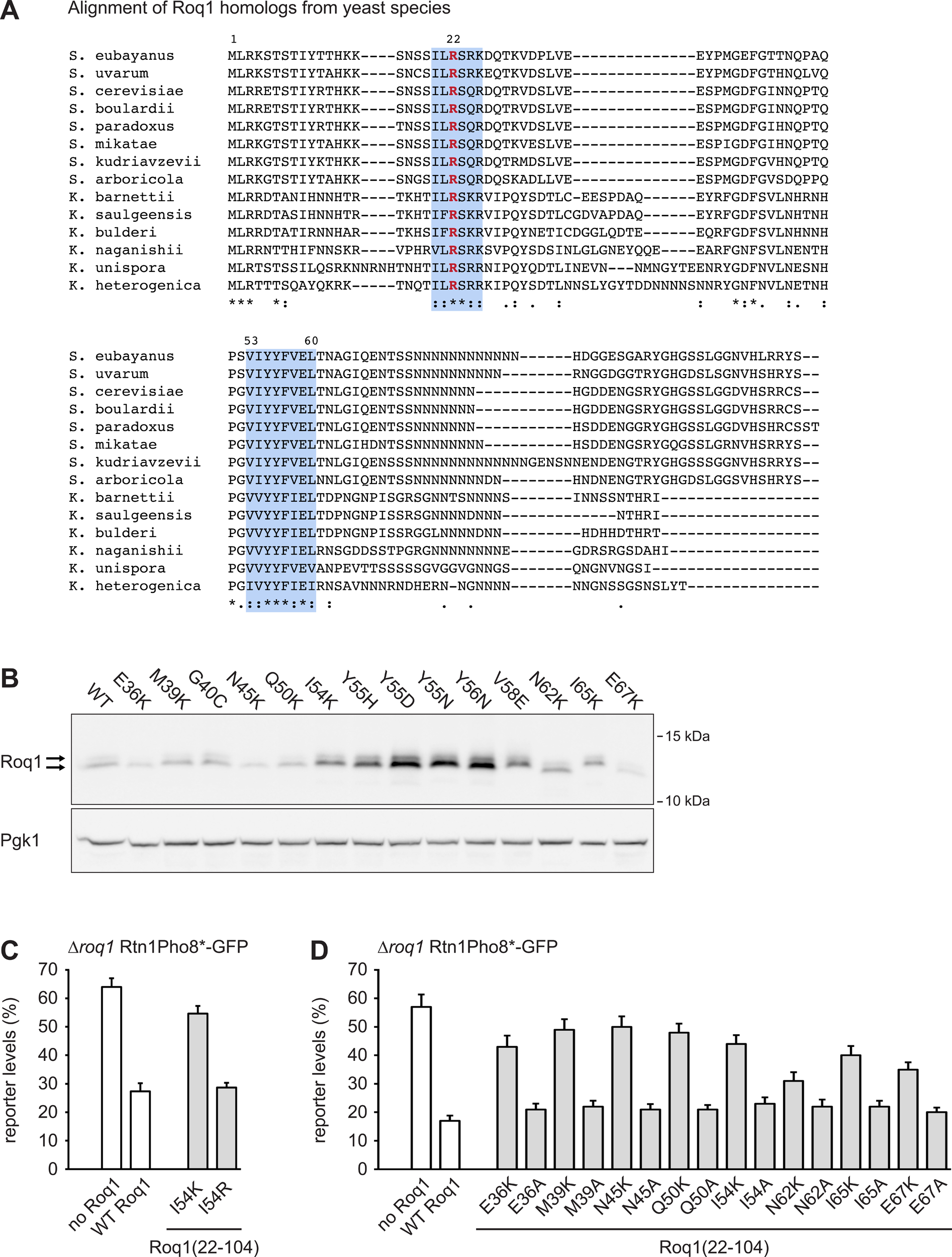
Sequence alignment of Roq1 homologs and expression and SHRED activity of Roq1 point mutants. A Multiple sequence alignment of Roq1 homologs from fourteen yeast species. The numbering of the residues corresponds to the *S. cerevisiae* sequence. * = fully conserved residue; : = strongly conserved residue; . = weakly conserved residue. B Western blot of HA tag and Pgk1 from *roq1* mutant strains expressing ubiquitin-Roq1(22-104)-HA(74) variants under the control of the strong *GPD* promoter. Roq1(22-104) naturally runs as a double band. The nature of the slower migrating band is unknown. Pgk1 served as a loading control. C, D Cellular levels of the SHRED reporter Rtn1-Pho8*-GFP after tunicamycin treatment for 5 h relative to the levels in untreated cells, as measured by flow cytometry. The *roq1* mutant cells contained an empty plasmid (no Roq1), a plasmid encoding wild-type Roq1 (WT Roq1) or plasmids encoding variants of ubiquitin-Roq1(22-104). Ubiquitin-Roq1 fusions are processed by cells to yield Roq1(22-104). Bars are the mean ± SEM; n = 3 biological replicates.

**Figure S4.**
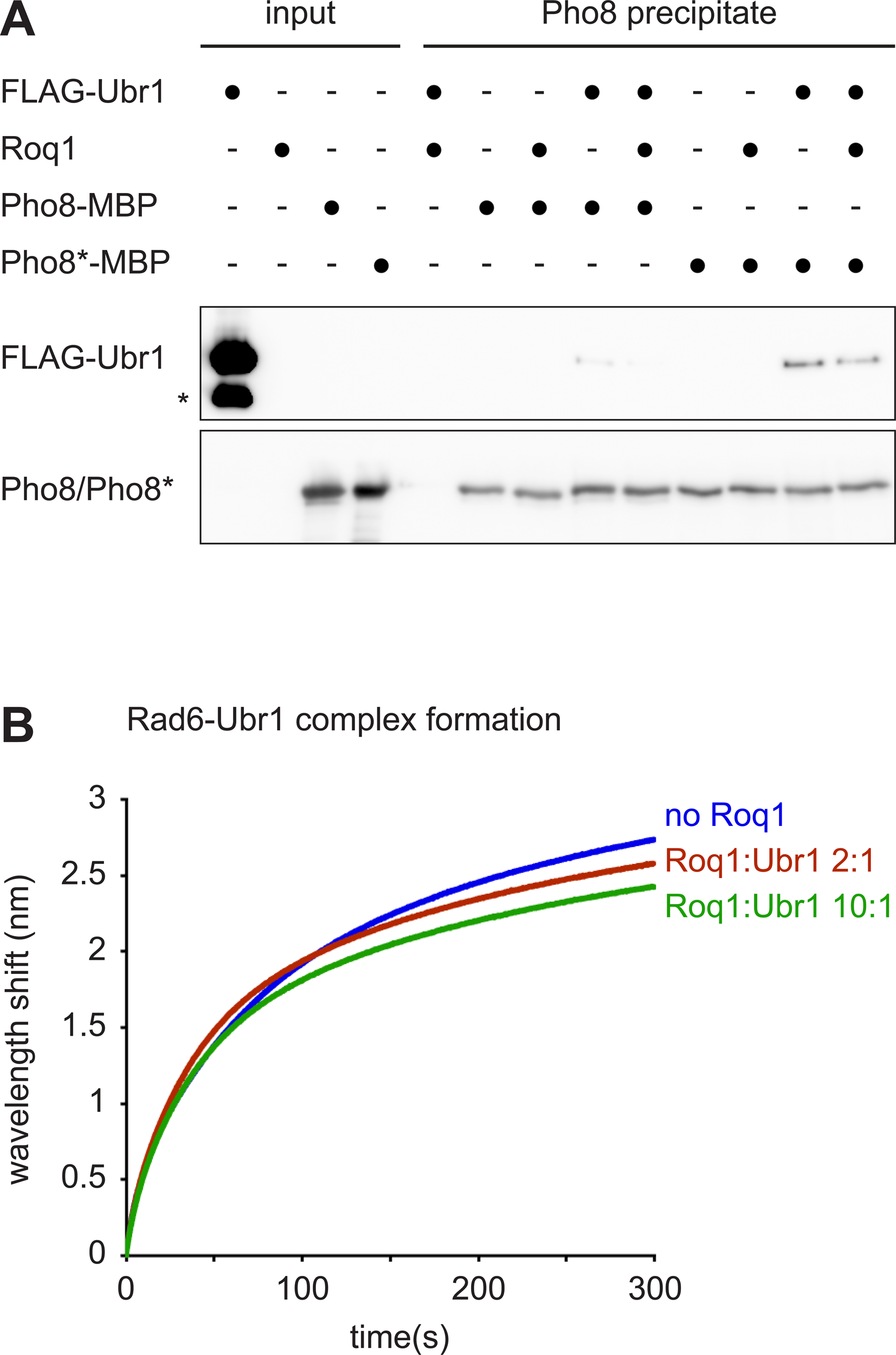
Roq1 does not enhance recognition of Pho8* or recruitment of Rad6. A Western blots of FLAG tag and Pho8 from input and Pho8 precipitate of an in vitro pulldown assays with FLAG-Ubr1, Pho8/Pho8* fused to maltose binding protein (MBP), and Roq1(22-60)-HA as indicated. Pho8 or Pho8*-MBP were precipitated with amylose resin. The asterisk denotes truncated Ubr1 that arose during the expression and purification of FLAG-Ubr1. Biolayer interferometry of Rad6-Ubr1 complex formation with immobilized Rad6 and soluble Ubr1. Complex formation was tested in the absence of Roq1(22-104) and with Roq1(22-104):Ubr1 molar ratios of 2:1 or 10:1.

**Figure S5.**
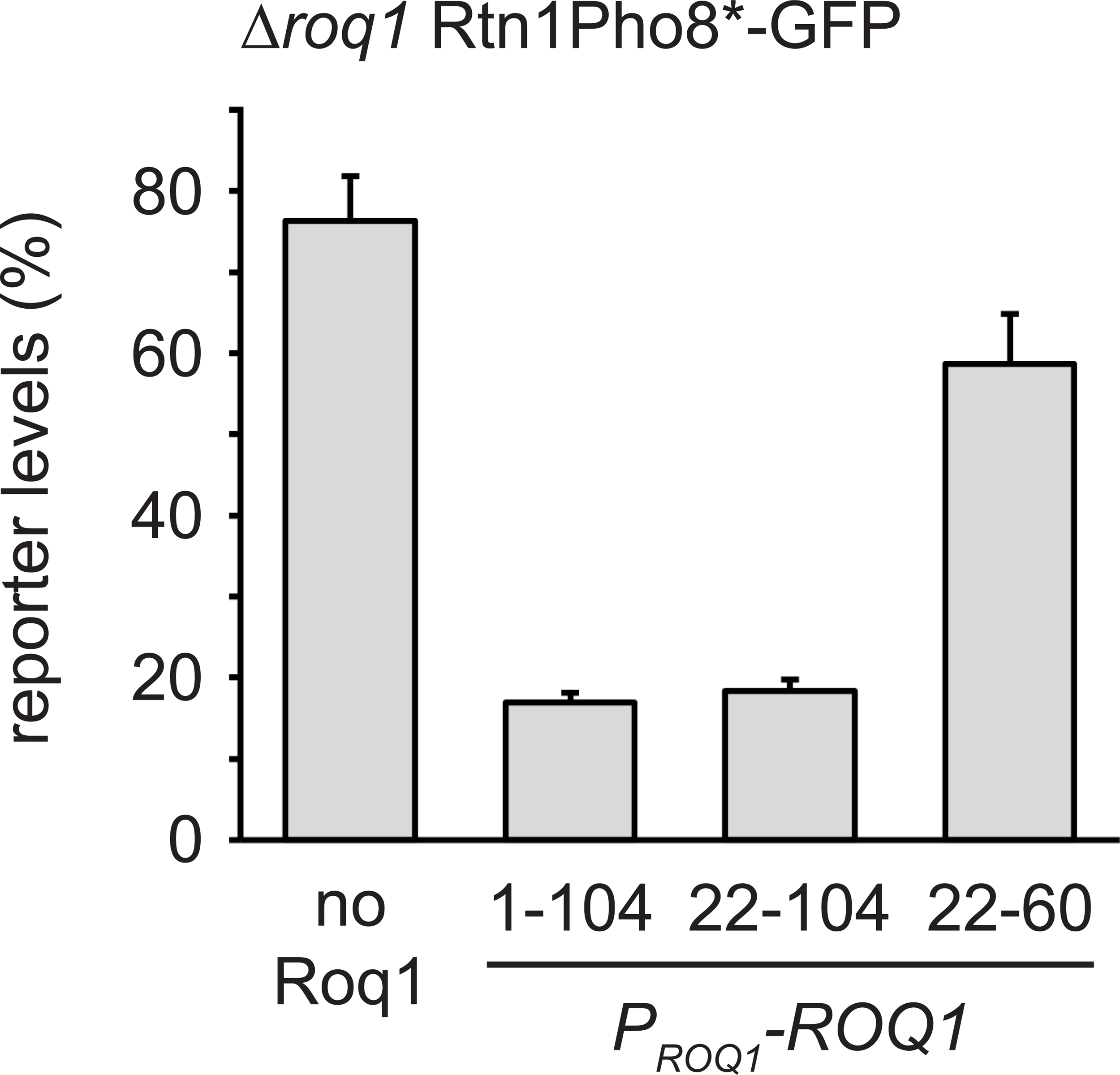
Roq1(22-60) shows reduced functionality in cells when expressed under the endogenous *ROQ1* promoter. Cellular levels of the SHRED reporter Rtn1-Pho8*-GFP after tunicamycin treatment for 5 h relative to levels in untreated cells, as measured by flow cytometry. The *roq1* mutant cells contained an empty plasmid (no Roq1) or expression plasmids for full-length Roq1, ubiquitin-Roq1(22-104) or ubiquitin-Roq1(22-60) that were controlled by the *ROQ1* promoter and integrated into the genome. Ubiquitin-Roq1 fusions are processed by cells to yield Roq1 starting with R22. Bars are the mean ± SEM; n = 3 biological replicates. P_ROQ1_, *ROQ1* promoter.

## References

Abramson J, Adler J, Dunger J, Evans R, Green T, Pritzel A, Ronneberger O, Willmore L, Ballard AJ, Bambrick J et al (2024) Accurate structure prediction of biomolecular interactions with AlphaFold 3. Nature, 630: 493–500

Andréasson C, Fiaux J, Rampelt H, Mayer MP, Bukau B (2008). Hsp110 is a nucleotide-activated exchange factor for Hsp70. J Biol Chem 283:8877–8884

Baker T, Varshavsky A (1991) Inhibition of the N-end rule pathway in living cells. Proc Natl Acad Sci U S A 88: 1090–1094

Buetow L, Huang DT (2016) Structural insights into the catalysis and regulation of E3 ubiquitin ligases. Nat Rev Mol Cell Biol 17: 626–642

Chakraborty S, Venkatramani R, Rao BJ, Asgeirsson B, Dandekar AM (2013) Protein structure quality assessment based on the distance profiles of consecutive backbone Cα atoms. F1000Res 2: 211

Chin JW, Martin AB, King DS, Wang L, Schultz PG (2002) Addition of a photocrosslinking amino acid to the genetic code of Escherichia coli. Proc Natl Acad Sci U S A 99: 11020–11024

Cowan AD, Ciulli A (2022) Driving E3 ligase substrate specificity for targeted protein degradation: lessons from nature and the laboratory. Annu Rev Biochem 91: 295–319

Cruz Walma DA, Chen Z, Bullock AN, Yamada KM (2022) Ubiquitin ligases: guardians of mammalian development. Nat Rev Mol Cell Biol 23: 350–367

Davey NE, Van Roey K, Weatheritt RJ, Toedt G, Uyar B, Altenberg B, Budd A, Diella F, Dinkel H, Gibson TJ (2012) Attributes of short linear motifs. Mol Biosyst 8: 268–281

Davey NE, Cyert MS, Moses AM (2015) Short linear motifs - ex nihilo evolution of protein regulation. Cell Commun Signal 13: 43

Deshaies RJ, Joazeiro CA (2009) RING domain E3 ubiquitin ligases. Annu Rev Biochem 78: 399–434

Du F, Navarro-Garcia F, Xia Z, Tasaki T, Varshavsky A (2002) Pairs of dipeptides synergistically activate the binding of substrate by ubiquitin ligase through dissociation of its autoinhibitory domain. Proc Natl Acad Sci U S A 99: 14110–14115

Eisele F, Wolf DH (2008) Degradation of misfolded protein in the cytoplasm is mediated by the ubiquitin ligase Ubr1. FEBS Lett 582: 4143–4146

Gottemukkala KV, Chrustowicz J, Sherpa D, Sepic S, Vu DT, Karayel Ö, Papadopoulou EC, Gross A, Schorpp K, von Gronau S et al (2024) Non-canonical substrate recognition by the human WDR26-CTLH E3 ligase regulates prodrug metabolism. Mol Cell 84: 1948–1963

Heck JW, Cheung SK, Hampton RY (2010) Cytoplasmic protein quality control degradation mediated by parallel actions of the E3 ubiquitin ligases Ubr1 and San1. Proc Natl Acad Sci U S A 107: 1106–1111

Holehouse AS, Kragelund BB (2024) The molecular basis for cellular function of intrinsically disordered protein regions. Nat Rev Mol Cell Biol 25: 187–211

Hwang CS, Shemorry A, Varshavsky A (2009) Two proteolytic pathways regulate DNA repair by cotargeting the Mgt1 alkylguanine transferase. Proc Natl Acad Sci U S A 106: 2142–2147

Jumper J, Evans R, Pritzel A, Green T, Figurnov M, Ronneberger O, Tunyasuvunakool K, Bates R, Žídek A, Potapenko A, et al (2021) Highly accurate protein structure prediction with AlphaFold. Nature 596: 583–589

Kelsall IR (2022) Non-lysine ubiquitylation: Doing things differently. Front Mol Biosci 9: 1008175

Kim JG, Shin HC, Seo T, Nawale L, Han G, Kim BY, Kim SJ, Cha-Molstad H (2021) Signaling pathways regulated by UBR box-containing E3 ligases. Int J Mol Sci 22: 8323

Kumar M, Michael S, Alvarado-Valverde J, Zeke A, Lazar T, Glavina J, Nagy-Kanta E, Donagh JM, Kalman ZE, Pascarelli S et al (2024) ELM-the eukaryotic linear motif resource-2024 update. Nucleic Acids Res 52: D442–D455

Kurgan L (2022) Resources for computational prediction of intrinsic disorder in proteins. Methods 204: 132–141

Mace PD, Day CL (2023) A massive machine regulates cell death. Science 379: 1093–1094

McShane E, Selbach M. (2022) Physiological functions of intracellular protein Degradation. Annu Rev Cell Dev Biol 38: 241–262

Mumberg D, Müller R, Funk M (1995) Yeast vectors for the controlled expression of heterologous proteins in different genetic backgrounds. Gene 156: 119–122

Oh JH, Hyun JY, Varshavsky A (2017) Control of Hsp90 chaperone and its clients by N-terminal acetylation and the N-end rule pathway. Proc Natl Acad Sci U S A 114: E4370–E4379

Pan M, Zheng Q, Wang T, Liang L, Mao J, Zuo C, Ding R, Ai H, Xie Y, Si D, Yu Y, Liu L, Zhao M (2021) Structural insights into Ubr1-mediated N-degron polyubiquitination. Nature 600: 334–338

Saita S, Nolte H, Fiedler KU, Kashkar H, Venne AS, Zahedi RP, Krüger M, Langer T (2017) PARL mediates Smac proteolytic maturation in mitochondria to promote apoptosis. Nat Cell Biol 19: 318–328

Schmidt R, Zahn R, Bukau B, Mogk A (2009) ClpS is the recognition component for Escherichia coli substrates of the N-end rule degradation pathway. Mol Microbiol 72: 506–517

Schmidt RM, Schessner JP, Borner GH, Schuck S (2019) The proteasome biogenesis regulator Rpn4 cooperates with the unfolded protein response to promote ER stress resistance. eLife 8: e43244

Schuck S, Gallagher CM, Walter P (2014) ER-phagy mediates selective degradation of endoplasmic reticulum independently of the core autophagy machinery. J Cell Sci 127: 4078–4088

Singer MA, Lindquist S (1998) Multiple effects of trehalose on protein folding in vitro and in vivo. Mol Cell 1: 639–648

Szoradi T, Schaeff K, Garcia-Rivera EM, Itzhak DN, Schmidt RM, Bircham PW, Leiss K, Diaz-Miyar J, Chen VK, Muzzey D, Borner GHH, Schuck S (2018) SHRED is a regulatory cascade that reprograms Ubr1 substrate specificity for enhanced protein quality control during stress. Mol Cell 70: 1025–1037

Taxis C, Knop M (2006) System of centromeric, episomal, and integrative vectors based on drug resistance markers for Saccharomyces cerevisiae. Biotechniques 40: 73–78

Tsai JM, Nowak RP, Ebert BL, Fischer ES (2024). Targeted protein degradation: from mechanisms to clinic. Nat Rev Mol Cell Biol doi: 10.1038/s41580-024-00729-9

Turner GC, Du F, Varshavsky A (2000) Peptides accelerate their uptake by activating a ubiquitin-dependent proteolytic pathway. Nature 405: 579–583

Ungelenk S, Moayed F, Ho CT, Grousl T, Scharf A, Mashaghi A, Tans S, Mayer MP, Mogk A, Bukau B (2016) Small heat shock proteins sequester misfolding proteins in near-native conformation for cellular protection and efficient refolding. Nat Commun 7: 13673

Van Roey K, Uyar B, Weatheritt RJ, Dinkel H, Seiler M, Budd A, Gibson TJ, Davey NE (2014) Short linear motifs: ubiquitous and functionally diverse protein interaction modules directing cell regulation. Chem Rev 114: 6733–6778

Varadi M, Bertoni D, Magana P, Paramval U, Pidruchna I, Radhakrishnan M, Tsenkov M, Nair S, Mirdita M, Yeo J et al (2024) AlphaFold Protein Structure Database in 2024: providing structure coverage for over 214 million protein sequences. Nucleic Acids Res 52: D368–D375

Varshavsky A (2011) The N-end rule pathway and regulation by proteolysis. Protein Sci 20: 1298–1345

Vittal V, Stewart MD, Brzovic PS, Klevit RE (2015) Regulating the regulators: recent revelations in the control of E3 ubiquitin ligases. J Biol Chem 290: 21244–21251

Wang L, Liu C, Yang B, Zhang H, Jiao J, Zhang R, Liu S, Xiao S, Chen Y, Liu B et al (2022) SARS-CoV-2 ORF10 impairs cilia by enhancing CUL2ZYG11B activity. J Cell Biol 221: e202108015

Wang B, Cao S, Zheng N (2024). Emerging strategies for prospective discovery of molecular glue degraders. Curr Opin Struct Biol 86: 102811

Xia Z, Webster A, Du F, Piatkov K, Ghislain M, Varshavsky A (2008) Substrate-binding sites of UBR1, the ubiquitin ligase of the N-end rule pathway. J Biol Chem 283: 24011–24028

Xie Y, Varshavsky A (1999) The E2-E3 interaction in the N-end rule pathway: the RING-H2 finger of E3 is required for the synthesis of multiubiquitin chain. EMBO J 18: 6832–6844

Young TS, Ahmad I, Yin JA, Schultz PG (2010) An enhanced system for unnatural amino acid mutagenesis in E. coli. J Mol Biol 395: 361–374

Zhang Y, Lin S, Peng J, Liang X, Yang Q, Bai X, Li Y, Li J, Dong W, Wang Y, et al (2022) Amelioration of hepatic steatosis by dietary essential amino acid-induced ubiquitination. Mol Cell 82: 1528–1542

Zheng N, Shabek N (2017) Ubiquitin ligases: structure, function, and regulation. Annu Rev Biochem 86: 129–157

Zhu K, Song L, Wang L, Hua L, Luo Z, Wang T, Qin B, Yuan S, Gao X, Mi W, Cui S. SARS-CoV-2 ORF10 hijacking ubiquitination machinery reveals potential unique drug targeting sites. Acta Pharm Sin B doi: 10.1016/j.apsb.2024.05.018

